# Determining mesoscale chromatin structure parameters from spatially correlated cleavage data using a coarse-grained oligonucleosome model

**DOI:** 10.1101/2024.07.28.605011

**Authors:** Ariana Brenner Clerkin, Nicole Pagane, Devany W. West, Andrew J. Spakowitz, Viviana I. Risca

## Abstract

The three-dimensional structure of chromatin has emerged as an important feature of eukaryotic gene regulation. Recent technological advances in DNA sequencing-based assays have revealed locus- and chromatin state-specific structural patterns at the length scale of a few nucleosomes (∼1 kb). However, interpreting these data sets remains challenging. Radiation-induced correlated cleavage of chromatin (RICC-seq) is one such chromatin structure assay that maps DNA-DNA-contacts at base pair resolution by sequencing single-stranded DNA fragments released from irradiated cells. Here, we develop a flexible modeling and simulation framework to enable the interpretation of RICC-seq data in terms of oligonucleosome structure ensembles. Nucleosomes are modeled as rigid bodies with excluded volume and adjustable DNA wrapping, connected by linker DNA modeled as a worm-like chain. We validate the model’s parameters against cryo-electron microscopy and sedimentation data. Our results show that RICC-seq is sensitive to nucleosome spacing, nucleosomal DNA wrapping, and the strength of inter-nucleosome interactions. We show that nucleosome repeat lengths consistent with orthogonal assays can be extracted from experimental RICC-seq data using a 1D convolutional neural net trained on RICC-seq signal predicted from simulated ensembles. We thus provide a suite of analysis tools that add quantitative structural interpretability to RICC-seq experiments.

## INTRODUCTION

Chromatin architecture regulates access to genomic DNA, playing a fundamental role in the control of gene expression and genome integrity. Nucleosomes, the fundamental unit of chromatin, are composed of 146-147 bp of DNA wrapped ∼1.7 times around a core of histone proteins (1, 2). The key geometric features of oligonucleosomes include: 1) the number of DNA base pairs wrapped around the nucleosome core; 2) nucleosome spacing, typically represented as the nucleosome repeat length (NRL); and 3) the stacking interactions between proximate nucleosomes. The number of wrapped base pairs can vary as nucleosomal DNA ‘breathes’ away from its core histone contacts. This breathing changes the angles at which DNA enters and exits the nucleosome core, thus affecting oligonucleosome geometry. Adjacent nucleosomes are connected by ∼10-75 bp of linker DNA (3, 4). Linker DNA length affects the relative orientation of neighboring nucleosomes—the addition of one base pair to a DNA linker changes the relative orientation of adjacent nucleosomes by approximately 34° due to the inherent helical twist of DNA (5, 6). Finally, oligonucleosome conformations are regulated by stacking interactions between proximate nucleosomes—mediated in large part by histone tails interacting with the histone core surface and the nucleosomal DNA (7, 8).

Oligonucleosome geometry plays an important role in determining the physical and functional properties of chromatin (9–15). For example, chromatin fibers with linker DNA lengths of 10n are more compact than those with 10n+5 linkers (16–20). Recent *in silico* work also shows that changes in nucleosomal DNA wrapping affect the propensity of oligonucleosomes to undergo phase separation (21). Inter-nucleosome interactions, modulated by histone modifications, themselves modulate chromatin fiber folding and structure (22–31). Finally, chromatin compaction and modification have been linked with transcriptional repression and modulation of the activity of chromatin-modifying complexes at the genome-wide level (32–35).

Locus-specific changes in chromatin structure at the length scale of regulatory DNA elements (∼200 bp to several kb) are likely to impact how macromolecular complexes interact with nucleosomes to drive DNA-based processes such as transcription, DNA replication, and DNA repair. However, the structure of chromatin on this length scale, the meso-scale, remains poorly understood. *In vitro,* chromatin formed from regular nucleosome arrays adopts 30-nm diameter fibers (36–38), which can be seen in rare instances in cells (39). In most cell contexts, however, advances in microscopy have shown that meso-scale chromatin organization is extremely heterogeneous and does not conform to the 30-nm fiber model (33, 40–45).

Current experimental methods are still limited in their ability to provide information about mesoscale chromatin structure. DNA sequencing-based methods like Micro-C (46–48) its variants (49, 50), and RICC-seq (51) are complementary to microscopy and produce data that are easy to integrate with chromatin features such as maps of histone modifications or transcriptional activity. Micro-C and Hi-CO have begun to identify tetranucleosomal chromatin folding motifs (52), that may be relevant to gene function. More work is needed to understand the relationship between chromatin’s mesoscale folding motifs and its epigenetic (post-translational modification) or functional states. Because these methods are based on enzymatic cleavage of DNA and crosslinking, it can be difficult to disentangle the three-dimensional contact signal from confounding enzymatic over-digestion and non-structural crosslinking biases (48).

RICC-seq, or radiation-induced correlated cleavage of chromatin with sequencing, can provide base-pair and nanometer-scale DNA-DNA contact information in intact cells (aggregated over defined genomic regions) without the need for enzymatic cleavage and crosslinking, and is thus methodologically orthogonal to Micro-C-like methods. As a relatively recently developed method, however, it lacks adequate tools for mechanistic interpretation of its results. Previously, RICC-seq signal was interpreted using 30-nanometer (nm) fiber structure models that assume regular helical morphologies (19, 51), which are unphysiological. Furthermore, RICC-seq has been used to build DNA fragment length distribution (FLD) histograms over aggregated loci of interest, usually as a function of epigenetic state. These FLDs do not provide enough information to fully specify a chromatin structure, but nevertheless are likely to contain important information about the average conformation of the chromatin fiber in regions of interest. The aim of this study was to develop flexible modeling tools to extract this structural information from RICC-seq FLDs.

The large molecular weight of oligonucleosomes makes them difficult to model at atomic or near-atomic resolution. Small oligonucleosome units have been modeled with minimal coarse-graining that retains chemical specificity (53). However, simulations with sufficient nucleosomes to capture the compaction and mechanical behavior of the chromatin fiber require additional coarse-graining. Most chromatin coarse-graining efforts have used rigid-body nucleosomes connected by semi-flexible DNA linkers discretized by beads (16, 54–58). The interactions between these nucleosomes have been modeled using coarse-grained electrostatics (21, 54–56) or using empirically determined or phenomenological effective potentials (16). Some recently developed multi-scale models also employ multiple levels of coarse-graining that validate against higher-resolution models (21, 56), and take locus-specific experimental data into account (59, 60). To interpret RICC-seq results, we need to understand how nucleosome spacing, nucleosomal DNA wrapping, and inter-nucleosome interaction values are reflected in the ssDNA fragment length distributions produced by radiation-induced correlated cleavage. We therefore built a model that was simple, customizable and efficient enough to explore and build oligonucleosome structure ensembles for a broad range of values for these geometric features.

Here, we adapt the kinked twistable wormlike chain (kinked tWLC) model—a Monte Carlo (MC) simulation framework that has previously been used to examine heterochromatin self-organization and compaction using longer length-scale chromosome simulations (10, 58, 61, 62)—for mesoscale chromatin simulations. This new model, which we call “meso-wlc,” has four major elements of added functionality. First, we explicitly model both nucleosomes and linker DNA segments as solid polyhedra with dimensions based on the mononucleosome and B-DNA crystal structures, respectively (63, 64). We identify nucleosome-nucleosome, nucleosome-DNA, and DNA-DNA steric clashes and reject MC steps that propose steric clashes. Second, we represent each DNA linker as multiple beads to capture more conformational detail than previous kinked tWLC models built with one bead per nucleosome (10, 58, 61, 62). Third, we implement an inter-nucleosome attraction potential to account for electrostatic interactions between nearby nucleosomes. The nucleosome orientation-dependent shape of this potential was previously determined *in silico* using the 3SPN-AICG model (28). Our model captures the face-face stacking, face-side interaction, and side-side interaction propensities of nucleosomes using the shape of this potential. Fourth, we calibrate the degree to which DNA wraps around the nucleosome to be consistent with experimentally observed entry-exit DNA angles in mono-nucleosomes.

To apply meso-wlc to RICC-seq, we generate ensembles of chromatin structures for a variety of nucleosome spacings, nucleosomal DNA wrapping values, and inter-nucleosome interaction strengths. We use these ensembles, combined with a simple distance-based model, to predict FLDs generated by spatially correlated DNA cleavage. These FLDs then serve as a training set for a neural network that allows us to extract chromatin structure parameters from both synthetic RICC-seq data generated on different, heterogeneous nucleosome arrays, and experimental RICC-seq data. We anticipate that this modeling framework will significantly improve the interpretability and impact of RICC-seq experiments.

## MATERIALS AND METHODS

### Coarse-grained mesoscale chromatin model

#### Coarse-grained nucleosome representation

Neighboring nucleosomes are connected by DNA segments (linker DNA) modeled as discrete, stretchable, shearable, twistable worm-like chains as previously implemented in kinked tWLC (10, 58, 61, 62) (See Figure 1A). The boundaries of each linker DNA segment are referred to as a “bead.” Its coordinates and orientation vectors are outputs of meso-wlc.

**Figure 1:**
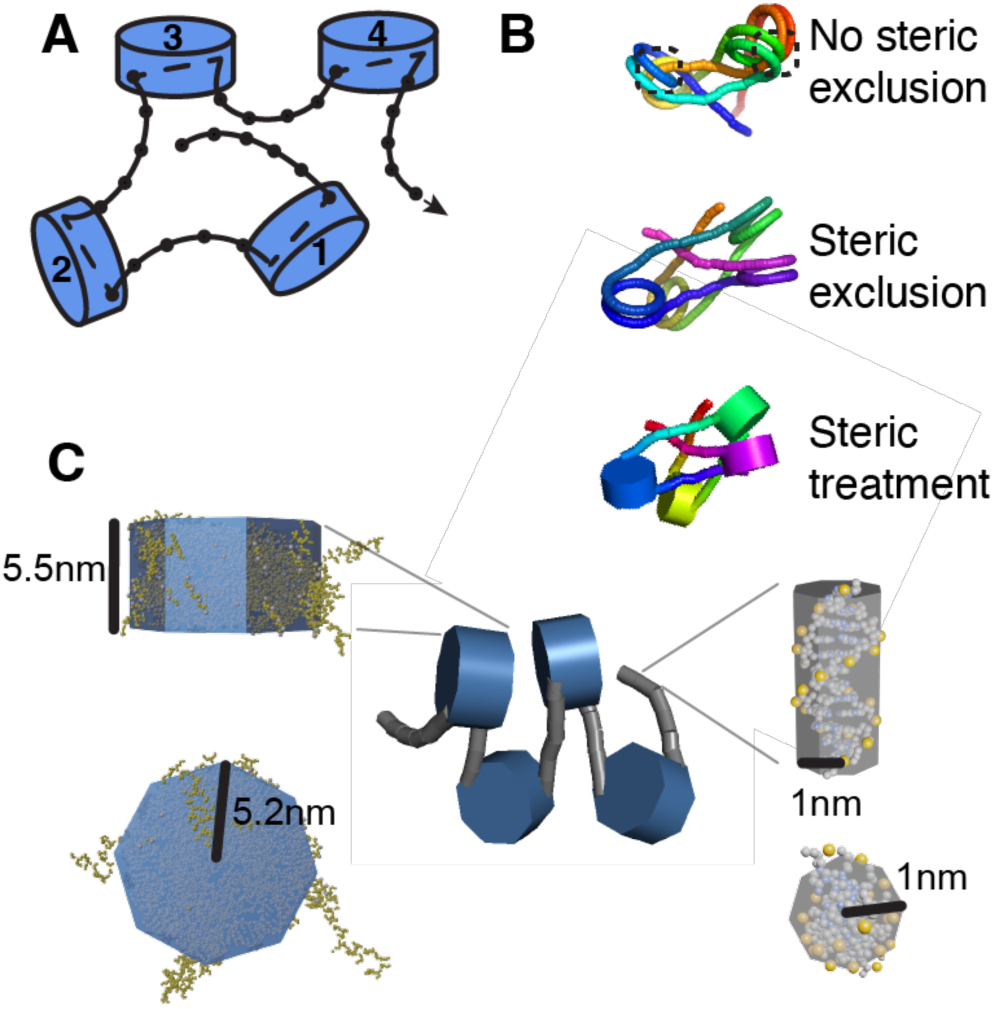
Coarse-grained representation of chromatin fibers. (**A**) **Schematic representation of coarse-grained modeling procedure**. Each linker DNA segment is modeled as a discrete, stretchable, shearable, twistable worm-like chain. Beads are represented by black circles. Nucleosomes are treated as rigid kinks (*10*). (**B**) **Steric collisions were detected and rejected using the GJK algorithm** (*65*). (Top) The depicted fiber is from a simulation without steric collision detection. Steric collisions are emphasized with black dashed boxes. (Middle) The depicted fiber is generated from a simulation with steric detection turned on. (Bottom) The depicted fiber is the same as the above but illustrates how each linker DNA bead and each nucleosome is modeled as an octagonal prism. The octagonal prisms’ vertices are input to the GJK algorithm to detect and reject collisions. (**C**) **Nucleosome core particles and DNA segments between beads are treated as octagonal prisms for steric collision detection**. (Left) The nucleosome core particle’s volume is modeled as an octagonal prism with radius 5.2 nm and height 5.5 nm, shown in blue. A mononucleosome crystal structure (PDB 1KX5) is shown docked inside the octagonal prism to demonstrate that modeling the nucleosome core particle as an octagonal prism with these dimensions is a reasonable approximation (*63*). Carbon atoms are shown in yellow. Phosphorous atoms are in gray. Atoms are scaled according to their Van der Waals radii. (Middle) A meso-wlc snapshot is shown with steric treatment illustrated. (Right) DNA between two computational beads is modeled as an octagonal prism with radius 1 nm and a height that scales with the number of base pairs between beads. The crystal structure of DNA (PDB 1BNA) is shown docked inside of an octagonal prism of radius 1 nm to demonstrate that modeling the linker DNA as an octagonal prism with radius 1 nm is a reasonable approximation (*64*). Carbon atoms are in gray, phosphorous are in yellow, and nitrogen are in blue.

Linker DNA segments are separated by a rigid kink, which comprises both a translational and rotational transformation, that accounts for the change in direction of the DNA chain as it goes through the nucleosome core particle (NCP), as previously implemented (10). The number of linker DNA beads in the system is given in Equation 1.

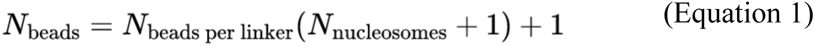

For steric collision detection (Figure 1B), the NCP is modeled as a rigid body with a regular octagonal prism shape with height 5.5nm (H_nuc_) and radius 5.2nm (R_nuc_). These parameters were obtained from the PDB 1KX5 nucleosome crystal structure (63) as shown in Figure 1C. The DNA between beads is also modeled as an octagonal prism with radius 1 nm (R_DNA_) and a height that scales according to the discretization. That is, H_DNA_ = *δz* where *δ* is the number of base pairs per bead and *z* is the distance between base pairs (See Supplementary Table S1 for parameter values).

#### Implementation of inter-nucleosome interactions in meso-wlc to enable mesoscale chromatin modeling

In order to model chromatin at the scale of 10^2^-10^3^ bp, where inter-nucleosome interactions and nucleosome geometries are important, we adjusted the kinked tWLC model to account for (1) nucleosome-nucleosome, nucleosome-linker DNA, and linker DNA-linker DNA steric clashes and (2) the attractive potential between nucleosomes.

After each Monte Carlo step, the Gilbert-Johnson-Keerthi (GJK) algorithm was used to detect steric clashes between objects modeled as octagonal prisms (See SI Text for more information) (65, 66). Fiber conformations with a steric clash were rejected (Figure 1B). This hard-shell potential and the GJK algorithm were chosen to maximize computational efficiency while capturing essential geometric features of chromatin.

An inter-nucleosome potential was also implemented to capture the energetically favorable interaction between the basic H4 tails and the acidic patch on H2A/H2B (67). We used the shape of the potential that the 3SPN-AICG model previously characterized to define inter-nucleosome interactions in meso-wlc as a function of distance and orientation (Supplementary Figure S1, Supplementary Figure S2) (28).

#### Parameter value estimation and model validation

Previously, a rigid “kink” was introduced to the tWLC model to account for DNA wrapping around the nucleosome core particle (kinked tWLC) (10). Entry and exit sites were determined by fitting 147 points, representing base pairs, around a nucleosome and propagating their twist and orientation vectors (10). The orientation vectors of these 147 base pairs account for the twist of B-form DNA with 10.5 base pairs per helical turn (6, 10).

The last helical turn of DNA on both the entry and exit side of the nucleosome does not wrap tightly around the nucleosome core particle (2). Therefore, modeling all 147 base pairs as a perfect helix results in over-wrapped structures as shown in Figure 2A. We computed the entry-exit angle for simulated mono-nucleosomes (Supplementary Figure S3) and compared the entry-exit angles from meso-wlc structures to entry-exit angles from experimental cryo-EM data to derive a more realistic nucleosome wrapping parameter of 127 bp (Figure 2A, Supplementary Figure S4) (68, 69).

**Figure 2:**
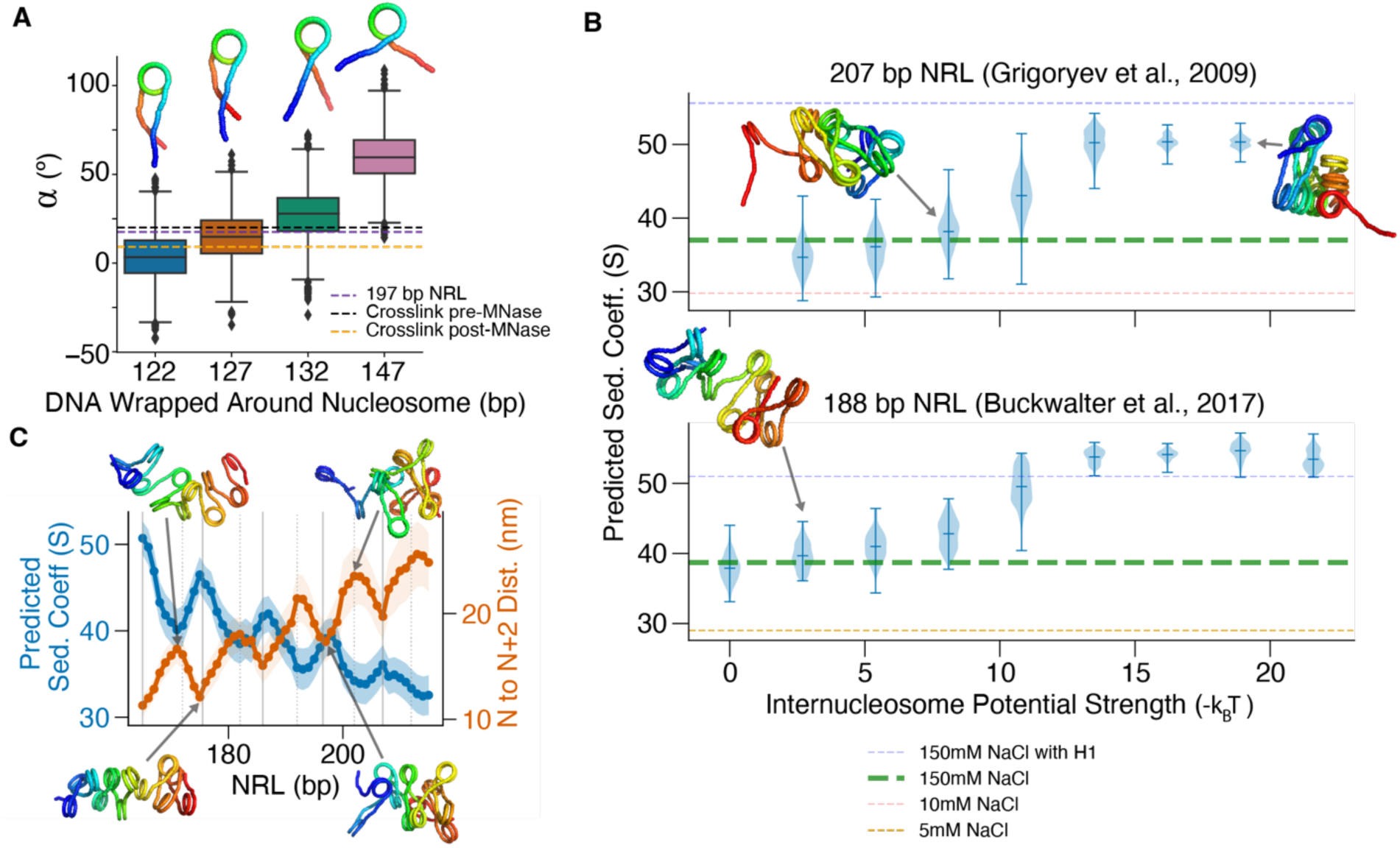
Meso-wlc parameters informed by experimental data yield fibers that demonstrate expected compaction trends. (**A**) A nucleosome wrapping parameter of 127 bp yields entry-exit angles consistent with Cryo-EM data of mono-nucleosomes, shown as a purple dashed line (*69*). Additional experimental data is shown as black and yellow dashed lines, See SI for additional information about these experimental conditions (*68*). Data is summarized over 15 trials and 191 snapshots are evaluated per trial. Each boxplot therefore represents 2,865 snapshots (See Supplementary Table S3 for additional parameters). (**B**) Scaling the inter-nucleosome potential to -5.4 kBT yields appropriately compact fibers. Linker histone (H1) and monovalent cations are known to aid in chromatin compaction. As expected, experimental conditions with H1 yield high sedimentation coefficients (blue dashed line) and low salt conditions yield low sedimentation coefficients (pink and yellow dashed lines). Simulated structures with inter-nucleosome potentials between 0 kBT and -8.1 kBT exhibit reasonable compaction as compared to *in vitro* arrays. (Top) As demonstrated by the snapshots shown, larger sedimentation coefficients result from more compact structures. Data from (top) Grigoryev et al., 2009 (*71*) and (bottom) Buckwalter et al., 2017 (*72*) (See Supplementary Table S4 for additional parameters). (**C**) Fiber compaction shows a periodicity of 10.5 bp. For each NRL, both the N-N+2 distance and the predicted sedimentation coefficient are computed across 4,365 snapshots (291 snapshots per trial for 15 trials) (Supplementary Table S5). Shaded area depicts one standard deviation. Solid vertical lines are separated by 10.5 bp, starting at 165 bp. Dotted vertical lines denote 10n+5 linker lengths.

We use sedimentation assay data from chromatin arrays without histone tails (70) as orthogonal data to validate this wrapping parameter selection. We used simulations with no internucleosome attractive potential as the most closely matched simulation condition because the chromatin arrays used in this experiment lack histone tails, which typically mediate a large portion of, though not all, inter-nucleosome contacts (7, 8). This allows us to calibrate wrapping alone without having to calibrate wrapping and the inter-nucleosome potential simultaneously. Meso-wlc ensembles with a wrapping parameter of 127 bp have predicted sedimentation coefficients that are consistent with experimental sedimentation coefficients at physiological salt concentrations (Supplementary Figure S5) (70), reinforcing our choice of default wrapping parameter at 127 bp.

Starting with an already validated model for the shape of inter-nucleosome potentials (28), we scaled the absolute strength of inter-nucleosome interactions relative to the DNA polymer elastic energy in our model by calibrating against *in vitro* sedimentation data. *In vitro* force spectrometry measured a nucleosome-nucleosome pair potential with a minimum of -1.6 kcal/mol (-2.7k_B_T) (73). Alternatively, fiber stretching experiments indicate that a more reasonable value is -3.4k_B_T in physiological salt (74) or -14k_B_T in high magnesium conditions (where low magnesium conditions destabilized nucleosome stacking) (75). These experiments contextualized the physiologically relevant range over which we scaled the inter-nucleosome potential in meso-wlc. We used meso-wlc to generate several ensembles of chromatin structures with an NRL of 188 bp, each with a different inter-nucleosome potential strength (Supplementary Table S4). Simulated structures had predicted sedimentation coefficients comparable to those observed *in vitro* at physiological salt when we set the inter-nucleosome potential scaling factor to 0 to -2.7k_B_T (Figure 2B). For the longer NRL of 207 bp, an inter-nucleosome potential of -5.4k_B_T is optimal. Fibers with longer linker DNAs have more flexibility than those with short NRLs and therefore the impact of this scaling factor increases with NRL. A scaling of -5.4k_B_T yields results generally consistent with the experimental data and can be set as the default parameter in meso-wlc, but users can select a scaling parameter that is appropriate for their questions of interest.

#### Running simulations

The number of beads per linker DNA segment was selected so that each bead represented approximately 8 bp. Simulations were initialized with beads in an outstretched initial orientation that helps to avoid both initialization biases in structure sampling and initial structures with steric clashes. If a sample is initialized with a steric clash, the clash is assigned a penalty of 3 k_B_T. The Monte Carlo algorithm is then implemented. Monte Carlo (MC) moves consist of crank shaft, translation, pivot, and single bead rotation moves (58). Each MC step consists of the quantity and type of move described in Supplementary Table S2. These moves were implemented in a previous version of kinked tWLC (58). If the conformation at the end of a step contains a new steric clash, the step is rejected. Otherwise, each MC step is accepted or rejected according to the Metropolis-Hastings criteria (76, 77). The number of steps per snapshot scales with the number of beads in the system and is reported for each simulation (Supplementary Table S3 - Supplementary Table S7).

The first 10 snapshots for each simulation were discarded to allow fiber equilibration. If energies did not equilibrate after 10 snapshots, a burn-in period of 50 snapshots was used. After this burn-in period, the distance between the center of the first and center of the last nucleosome was computed (the end-to-end distance). We then checked several simulations to ensure that the end-to-end distances were not autocorrelated between snapshots (See SI for more information, Supplementary Figure S6).

Each simulated structure had 12 nucleosomes and 127 bp wrapped around the nucleosome core particle, unless otherwise specified. Specific parameters for each simulation are given in Supplementary Information (Supplementary Table S3 - Supplementary Table S7).

#### Heterogeneous linker simulations

For the heterogeneous linker simulations, all linker DNA segments were represented by 8 beads. Each linker DNA length was drawn from a distribution of interest. Otherwise, simulations were run the same as the simulations with constant linker lengths (See Supplementary Table S8 for additional simulation details).

### In silico Radiation-Induced spatially Correlated Cleavage (RICC) prediction model

#### Interpolation of phosphate backbone positions

The signal produced by RICC-seq is a distribution of single-stranded DNA fragment lengths measured at base-pair resolution (51). However, meso-wlc does not have base-pair resolution. Approximately 8 base pairs are represented by a single coarse-grained computational bead. Predicting fragment lengths from meso-wlc structures requires interpolating the phosphate backbone positions between coarse-grained beads. In Equation 2, we define the double helix of B-DNA as two right-handed helices (78). Equation 2 yields the locations of the two phosphate backbones (r_p+_ and r_p-_) from both the plus (+) and minus (-) strands as well as the base-pairing location (r_bp_) between them.

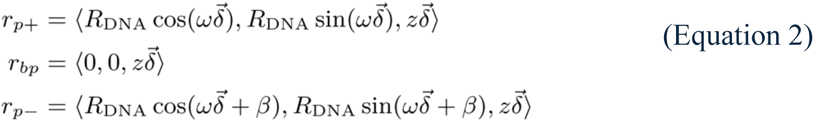

All constants (R_DNA_, ω, β, z) describing B-DNA double helix are given in Supplementary Table S1. The vector 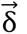 is the array {1… δ}, where δ is the number of base pairs per computational bead.

The base pair coordinates are interpolated between coarse-grained beads (Figure 3A, B). A three-sites-per-base-pair construction for our nucleosomal DNA is obtained by applying Equation 2 to the left-handed nucleosome superhelix (Figure 3B).

**Figure 3:**
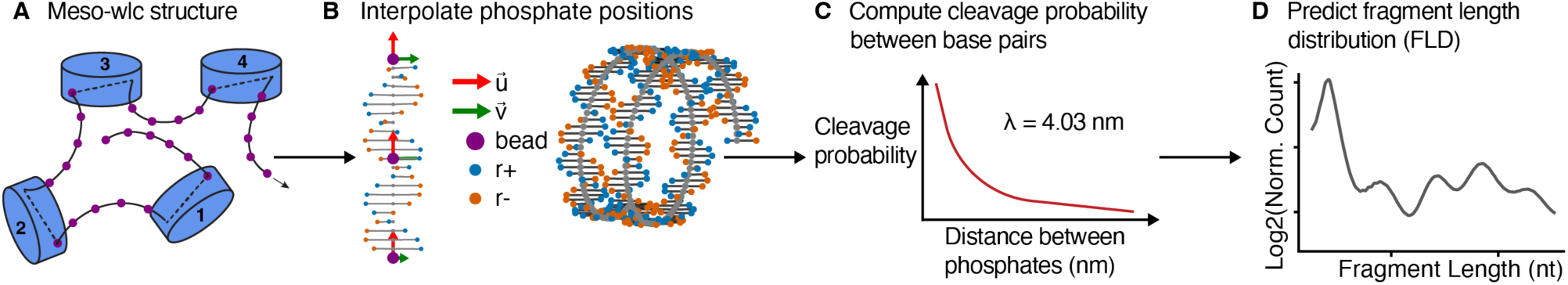
Phosphate backbone positions are interpolated between coarse-grained meso-wlc beads and used to predict fragment length distributions. (**A**) A schematic of a meso-wlc structure is shown with meso-wlc bead coordinates highlighted in purple. (**B**) Equation 2 is applied to interpolate DNA phosphate backbone positions. Each meso-wlc bead has a tangent vector 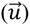 and an orientation vector 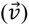. (**C**) Schematic of how distances between phosphates are passed through an exponential decay function with a rate constant (λ) of 4.03 nm (*51*). Cumulative cleavage likelihoods are referred to as counts. (**D**) Example of a predicted FLD. Using the counts and the 1D distance between phosphates (fragment lengths), we generate a distribution of the fragment lengths (See Materials and Methods: Fragment length distribution prediction for more information).

#### Fragment length distribution prediction

RICC-seq fragment counts around positioned nucleosomes were previously compared with the distances between pairs of DNA backbone phosphates in a mononucleosome crystal structure. An exponential fit of these data yielded that a single exponential with a length constant of 4.03 nm describes the relationship between 3D distance and relative cleavage probability (51) (Figure 3C). This relationship was used to construct the following *in silico* fragment length prediction model (Figure 3D):

First, a distance matrix (D) for each DNA strand is populated with the distances between each pair of interpolated phosphate backbone positions:

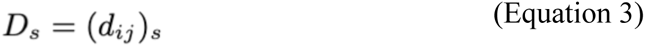

where *s* denotes either the plus strand or minus strand. The Euclidean distances (*d_ij_*) are computed between phosphate backbone positions in the same strand:

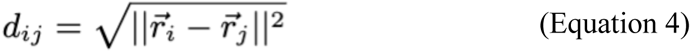

The probability of two nucleotides cleaving together given their positions (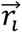 and (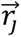) is given by:

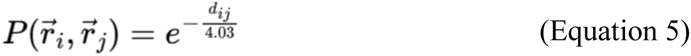

The normalized counts of fragment length *f* being observed in a simulated fiber of length *n* base pairs is given by:

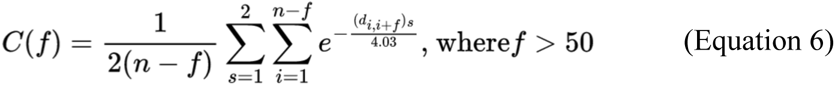

Fragment lengths below 50 bp are disregarded because fragments of these lengths are excluded during sequencing library preparation to remove adapter dimers. Short read sequencing has a bias against longer fragment lengths; especially, those of length >500 bp (79). The analysis in this manuscript evaluates only fragments <500 bp, but the length bias will likely result in the over-representation of longer fragments in the predicted fragment length distribution (FLD) compared to what would be expected from a sequencing experiment. This length bias can vary based on experimental conditions (79) and is not accounted for in this *in silico* model but can be accounted for when comparing simulated results to experimental results. That is, the length bias can be corrected in the experimental data as has been previously done (51). The process of generating a predicted FLD from a meso-wlc structure is illustrated in Figure 3.

### Convolutional neural nets trained on predicted fragment length distributions

We employed 1-dimensional convolutional neural nets (1D-CNN) to infer nucleosome repeat length, nucleosome wrapping, and inter-nucleosome interaction strength from predicted fragment length distributions. The predicted FLDs are max-normalized. Each 1D-CNN was trained with mean square error (MSE). The 1D-CNNs were comprised of 2 convolutional layers (each with a kernel size of 10 and a dilation size of 5), 2 max pooling layers, a dropout layer after the first pooling layer, and two linear layers. The models used leaky ReLU activation functions and were implemented using PyTorch v2.0.1 (80). The kernel parameters were selected to span 50 bp of DNA, which is less than one gyre of DNA wrapped around the nucleosome core particle. The dimensions of the output layer correspond to the number of parameters being predicted— a model that predicts only NRL has a 1-dimensional output layer whereas a model that predicts NRL and wrapping simultaneously has a 2-dimensional output layer. The number of training epochs varied per model and are reported in Supplementary Table S9. The models were trained with 66.67% of the data and the remaining 33.33% was used for testing.

### RICC-seq data processing

RICC-seq read pairs were aligned to hg19 and filtered for mapping quality and overlap with blacklisted regions as previously described (51). These reads were subset (81) based on intersection with ATAC-seq peaks called in the same cell line (51) and used to make an FLD histogram based on the aggregated open chromatin regions detected using ATAC-seq. This was compared against the genome-wide FLD.

### Computational resources

Simulations were run on an 8,400+ core HPC cluster running 64-bit Red Hat Linux 7 and using a variety of Intel processors, with one core per simulation trial.

## RESULTS

### Nucleosome spacing-dependent compaction

We verified that the compaction of the simulated chromatin ensembles as a function of NRL is consistent with previous observations. (16–18). We ran 51 simulations of 12-mer oligonucleosomes with NRLs ranging from 165 to 215 bp and 15 trials for each simulation, and evaluated 291 snapshots per trial (See Supplementary Table S5 for additional parameters). Our simulated ensembles indeed exhibit an NRL-dependent oscillation in the distance between a nucleosome (N) and its second-nearest nucleosome along the array (N+2) with a period of 10.5 base pairs, the helical turn of DNA (Figure 2C). The maxima of the oscillations are at linker lengths that are multiples of 10n+5 bp, which produce the least compact chromatin fibers; linker lengths of 10n bp are more compact than their 10n+5 counterparts. We also measured overall compaction of the oligonucleosomes via the predicted sedimentation coefficient and found a similar periodic dependence on the NRL (Figure 2C), with a period of 10.5 bp, consistent with the previously observed behavior.

### Nucleosome wrapping-dependent compaction

We then investigated the effect on chromatin compaction of nucleosome wrapping, which changes the entry-exit angle of nucleosomal DNA. Under physiological conditions, the number of base pairs wrapped around the NCP can be altered by thermal fluctuations, or ‘breathing’ of the nucleosomal DNA, especially for the first 10 bp wrapping on each side (2). We ran 23 simulations of a nucleosome array with 12 nucleosomes and an NRL of 177 bp with 117 to 147 bp of DNA wrapped around the nucleosome core particle. We ran 10 trials for each simulation and 291 snapshots were analyzed per simulation (See Supplementary Table S6, indices 17-39 for additional simulation parameters). We observe that nucleosome wrapping also affects the compaction of the chromatin fiber, as has been previously measured (5, 82–84). Using the predicted sedimentation coefficient as a measure of fiber compaction, we observed that the compaction of the chromatin fiber increases with nucleosome wrapping until reaching a plateau around 129 bp and decreasing slightly at 147 bp (Supplementary Figure S5). These results indicate that our chosen nucleosome wrapping parameter of 127 bp gives rise to a level of compaction consistent with experimental measurements (Supplementary Figure S5) (70).

### Nucleosome spacing-dependent and wrapping-dependent compaction are detectable by simulated RICC-seq signal

In order to use our chromatin ensembles to understand RICC-seq data, we predicted fragment length distributions for each meso-wlc snapshot using exponential-falloff of correlated cleavage frequency with distance (Figure 3, see Materials and Methods) (51). As previously observed for synthetic FLDs predicted from symmetric 30-nm fiber models (51), the shape of the FLD depends on the NRL with similar shapes approximately every 10 bp (Supplementary Figure S7). We also observed that RICC-seq fragment length distributions predicted from our simulated chromatin fiber ensembles are sensitive to nucleosome wrapping, suggesting that RICC-seq has the ability to measure this geometric parameter of chromatin. We ran 10 trials each of 16 simulations with an NRL of 185 bp and 117-147 bp wrapped around the nucleosome core (See Supplementary Table S6, indices 1-16 for additional parameters).

The location of the first peak in the fragment length distribution is consistent with the length of a ssDNA fragment arising from correlated breaks at an intra-nucleosome DNA gyre-to-gyre contact—approximately 80 nucleotides (nt), or one wrap of DNA around the nucleosome core (Figure 4A). This peak appears at the same location independent of nucleosome wrapping because the length of one nucleosome wrap depends only on the diameter of the nucleosome core. The second peak corresponds to a single nucleosome peak and the summit should correspond to where the entry and exit DNA linkers cross. Increased wrapping should correspond to a shorter peak center. As the nucleosome unwraps, the intersection between the entry and exit DNA should occur further away, corresponding to a longer fragment. This expected behavior is observed; wrapping is observed to be inversely correlated with the 2nd peak location (Figure 4B). The width of the distribution increases as wrapping decreases (Figure 4B). For an entry-exit angle (2α) of 90°, we would expect a narrow distribution, whereas the distribution should broaden as the entry-exit angle approaches 0°.

**Figure 4:**
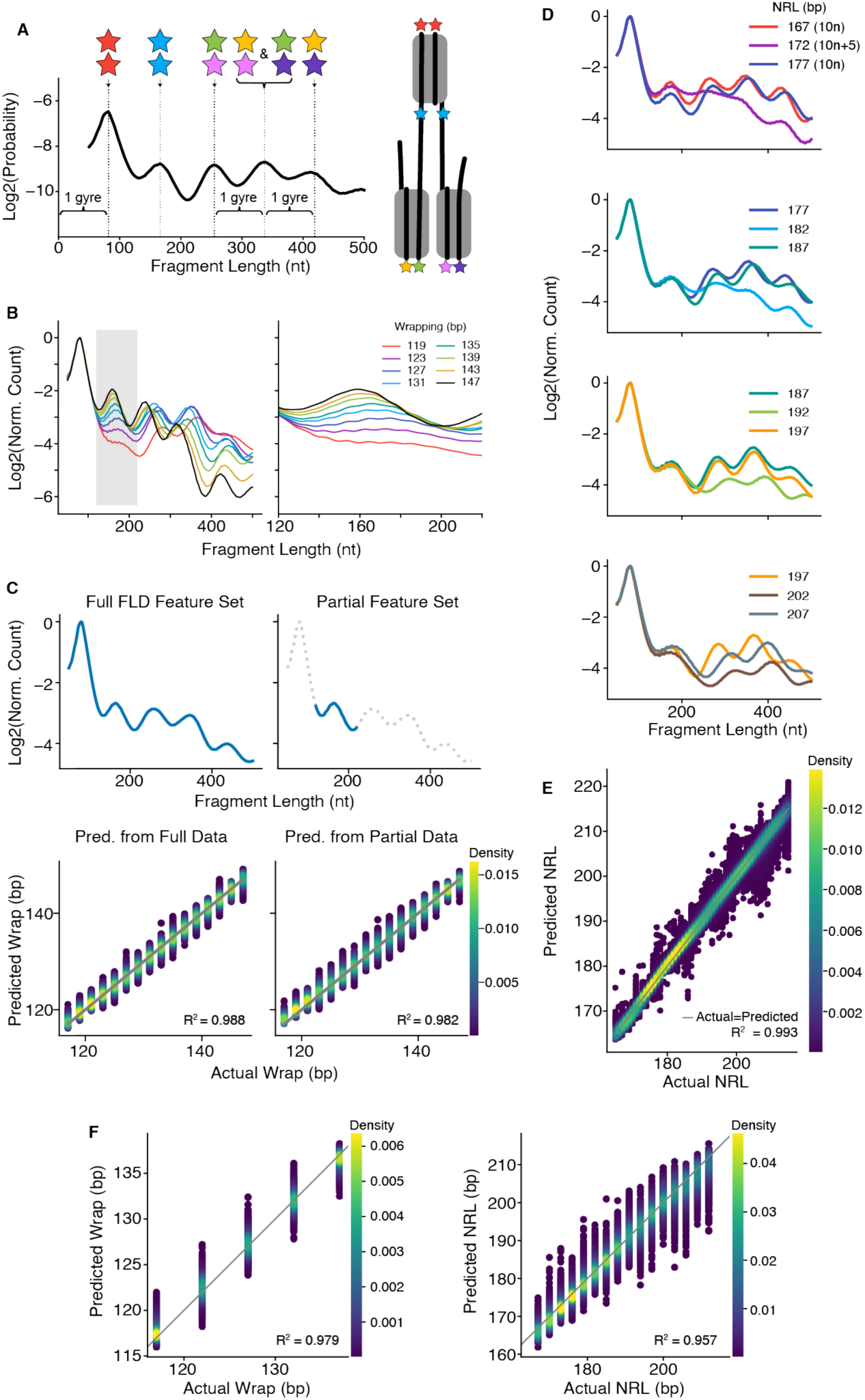
Nucleosome spacing and wrapping information is embedded in the FLDs. (**A**) Fragment length distribution from a simulation with 137 bp wrapping and 177 bp NRL is shown along with a cartoon illustrating from where the peaks originate. (**B**) (left) Predicted RICC-seq signal is sensitive to wrapping. As the wrapping parameter increases, the second peak shifts to the left and becomes more narrow. (right) A zoomed in view of the second peak is shown. Fragment length distributions are max-normalized before log transformation. (**C**) (left) The wrapping parameter is reliably inferred from the predicted RICC signal. The diagonal line shows where the x- and y-axis values are equivalent. (right) A model trained only on the data from the second peak performed well at predicting wrapping. Simulations were performed with a fixed NRL of 185 bp. (**D**) Predicted RICC-seq signal shows 10n+5 fibers (172, 182, 192, 202 NRLs) are less compact than both their 10n (167, 177, 187 NRLs) and 10n+10 (177, 187, 197, 207 NRLs) counterparts. The 3rd, 4th, and 5th peaks are less pronounced in 10n+5 fibers. (**E**) The NRL is reliably inferred from the predicted RICC signal. The diagonal line shows where the x- and y-axis values are equivalent. Simulations were performed with a fixed wrapping of 127 bp (**F**) The NRL and wrapping parameter are reliably inferred from predicted RICC signal simultaneously. Model is trained on simulations with NRLs ranging from 167 bp to 212 bp and wrapping parameters ranging from 117 to 137 bp.

The location of the 3rd peak also depends on nucleosomal DNA wrapping. The fragment lengths comprising the distribution’s third peak are consistent with a contact between the nearest DNA gyres of nucleosomes N and N+2 along the chromatin fiber (Figure 4A, green-pink contact). The fourth and fifth peaks occur at the same location relative to the third peak. The fourth peak is 1 gyre (82 nt) away from the third peak and the fifth peak is one gyre away from the fourth peak (Figure 4A, yellow-pink/green-purple, and yellow-purple, respectively).

#### Model 1: 1D-CNN trained on predicted FLDs from simulations with various nucleosomal DNA wrapping parameters

We analyzed the dependence of the simulated fragment length distributions on nucleosome wrapping at a constant NRL using a one-dimensional convolutional neural net (1D-CNN). We ran 10 trials with an NRL of 185 bp for each wrapping parameter. Each trial generated 201 snapshots, 10 were discarded to allow for equilibration, so 191 snapshots were analyzed for each trial (See Supplementary Table S6, indices 1-16 for additional simulation parameters). An FLD was predicted for each snapshot using the fragment length prediction model (see Materials and Methods). For each snapshot, the FLD was the feature set and the wrapping parameter was the target. We used 67% of the snapshots to train the model and 33% to test it. We trained the 1D-CNN model on this training set to predict the wrapping from the FLD. The regression performed with an R^2^ = 0.988 and RMSE = 1.01 bp demonstrating that, for constant NRL, nucleosomal DNA wrapping can be inferred from predicted RICC-seq signal (Figure 4C).

#### Model 2: 1D-CNN trained on ablated FLDs from simulations with various nucleosomal DNA wrapping parameters

We hypothesized that all the information about wrapping could be gleaned from the FLD data from the second peak, which we defined as the 100 nt region 120-219 nt (Figure 4B). To test this hypothesis, we ablated all of the FLD data outside of this fragment length range and trained a new 1D-CNN on only the data from the second peak of the FLD (Figure 4C). Consistent with our hypothesis, this model performed comparably to the model trained on the full FLD (R^2^=0.982, RMSE = 1.23 bp) (Figure 4C). This reveals that RICC-seq signal not only encodes information about nucleosome wrapping, but that this wrapping information can be inferred from the shape and location of the second FLD peak.

#### Model 3: 1D-CNN trained on predicted FLDs from simulation with various nucleosome spacings

The decreased compaction for chromatin fibers with 10n+5 bp linker lengths is also visible in predicted RICC fragment length distributions (Figure 4D). Simulations of ensembles of 10n and 10n+5 fibers reveal that 10n+5 fibers are generally more unfolded than their 10n counterparts, as evidenced by the less pronounced 3^rd^-5^th^ FLD peaks (Figure 4D). This indicates that the relative compaction is intrinsic to the geometric properties of the fiber and supports that these properties are experimentally detectable with RICC-seq.

We then used a 1D-CNN to analyze the effect on the simulated fragment length distributions of the NRL at constant wrapping. We trained the model on predicted FLDs from simulations with NRLs ranging from 165 bp-215 bp at a constant wrapping parameter of 127 bp. For each NRL, we performed 15 trials of each simulation (See Supplementary Table S5 for additional parameter details). An FLD was predicted for each snapshot. The regression performed with an R^2^ = 0.993 and RMSE = 1.19 bp. We conclude that, for constant wrapping, NRL can be inferred from predicted RICC-seq signal using this 1D-CNN approach (Figure 4E).

#### Model 4: 1D-CNN trained on predicted FLDs from simulations with various nucleosome spacing and nucleosome wrappings

We sought to investigate whether the 1D-CNN approach can simultaneously infer both NRL and nucleosomal DNA wrapping if it is trained on simulations in which both parameters are varied. The combined training and testing data had dimensions 69,840 by 451 where 69,840 snapshots were represented and 451 is the length of the FLD (50-500 bp). The output layer of the 1D-CNN had 2 dimensions, one corresponding to NRL and the other corresponding to wrapping. The 69,840 snapshots are from 5 DNA wrapping values spanning 117-137 bp and 16 NRLs spanning 167-212 bp (Supplementary Table S7). The model performed with an R^2^ = 0.957 and RMSE = 2.87 bp for NRL predictions and an R^2^ = 0.979 and RMSE = 1.03 bp for nucleosomal wrapping predictions (Figure 4F, Supplementary Table S9), indicating that both values can be simultaneously learned and predicted by the 1D-CNN.

### 1D-CNN can infer inter-nucleosome potential scaling from fragment length distribution

Histone modifications that change the inter-nucleosome potential are of epigenetic importance (27, 32). For example, acetylation of the H4 tail is associated with transcriptionally active chromatin and acetylation reduces the magnitude of the inter-nucleosome potential (22–26, 28–31). We therefore explored how the inter-nucleosome potential affects the RICC-seq signal, to determine if we can observe changes akin to epigenetic state differences with our simulations. Scaling up the inter-nucleosome potential increases the frequency of N-N+2 contacts as evidenced by the increased density in the 3^rd^-5^th^ predicted FLD peaks (Figure 5).

**Figure 5:**
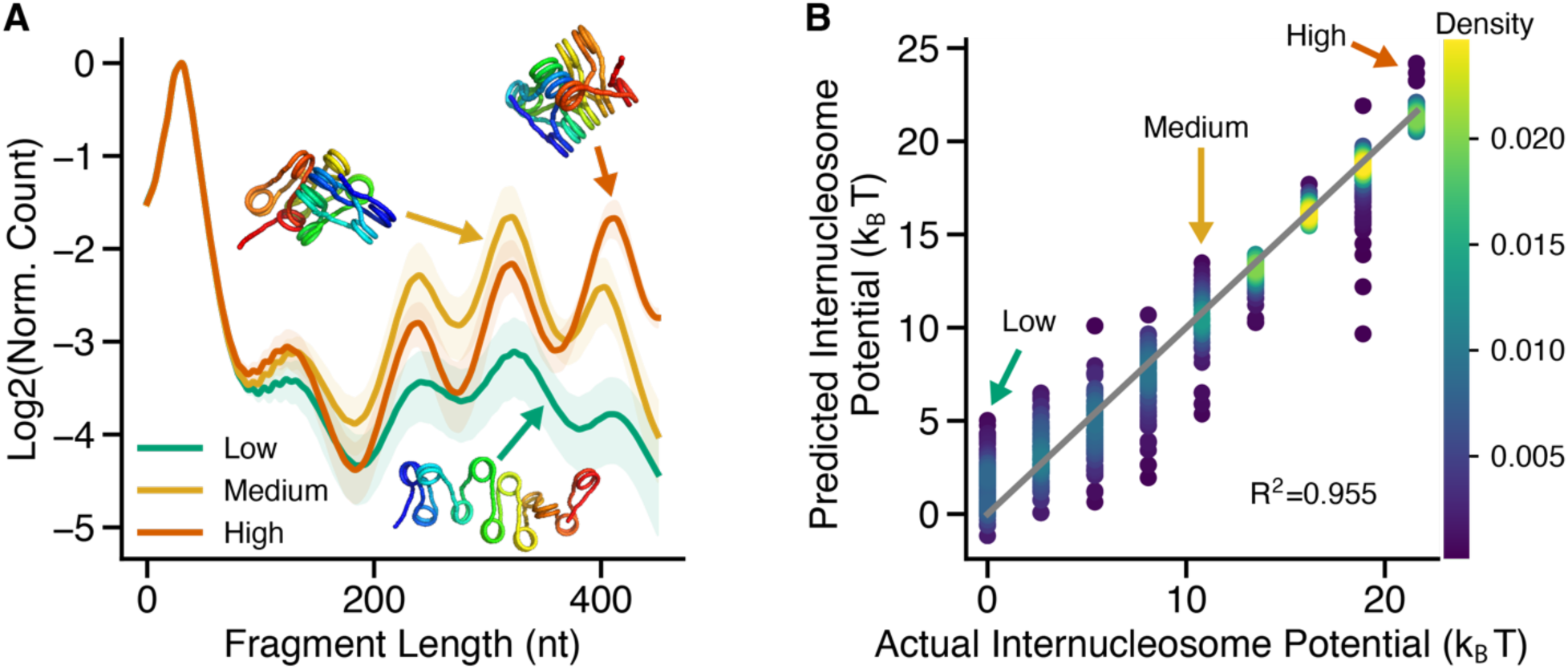
The inter-nucleosome potential scaling factor affects the predicted RICC-seq FLD. **(A**) As the inter-nucleosome potential is scaled up, more density in the 3rd-5th peaks is observed. This is expected because scaling up the inter-nucleosome potential facilitates N-N+2 interactions which comprise the 3rd-5th peaks. Low, Medium, and High potentials are 0 kBT, 10.8 kBT, and 21.6 kBT, respectively. Shaded regions represent 1 standard deviation. Representative snapshots are shown for each simulation. (**B**) 1D-CNN regression results. Simulations were performed with 188 bp NRL, 453 snapshots were evaluated per inter-nucleosome potential. Additional simulation parameter values can be found in Supplementary Table S4 (panel A: indices 8,12,16, panel B: indices 8-16).

#### Model 5: 1D-CNN trained on predicted FLDs from simulation with various inter-nucleosome potential magnitudes

Using the same neural net architecture described above, we trained a new neural net on predicted fragment length distributions from simulations of 12mers with a nucleosome repeat length of 188 bp. The model is able to reliably predict the inter-nucleosome potential with R^2^ = 0.955 and RMSE = 0.563 k_B_T (Figure 5). This demonstrates the principle that the relative strength of the inter-nucleosome potential is detectable in simulated RICC-seq signal.

### 1D-CNN trained on structural ensembles of homogeneous linker lengths performs well on ensembles with some linker length heterogeneity

The structural ensembles used to train Model 3 had homogeneous linker DNA lengths. However, physiological nucleosome spacing can be heterogeneous (85–88), and both experimental and computational studies have indicated that heterogeneity in nucleosome spacing can have profound effects on chromatin fiber conformations (10, 11, 13–15). SAMOSA-seq data from K562 cells indicated that 14-17 bp was a typical median absolute deviation in nucleosome spacing among regularly spaced nucleosome clusters (85). We therefore tested the extent to which heterogeneity in the simulated linker lengths affects the ability to predict average NRLs from simulated FLDs. We ran 6 simulations. In each simulation, the 13 linker DNA lengths were drawn from a distinct distribution (See Figure 6A and Supplementary Table S8 for additional simulation details). We then predicted FLDs from each snapshot and applied Model 3 (which was trained on chromatin structures with homogeneous linker lengths) to predict the NRLs from structures with linker length heterogeneity. We found that NRLs within 6 base pairs of the ground-truth average are predicted in structural ensembles with heterogenous linker lengths (Figure 6A) despite the fact that linker length heterogeneity appears to have a strong impact on the predicted FLD (Figure 6B). It would be challenging to run enough simulations with heterogenous linker lengths to cover the parameter space of possible NRL means and standard deviations to train the model on heterogeneous fibers. However, it is promising that this limited exploration nevertheless shows that a model trained on homogeneous linker lengths is robust to some heterogeneity in linker lengths.

**Figure 6:**
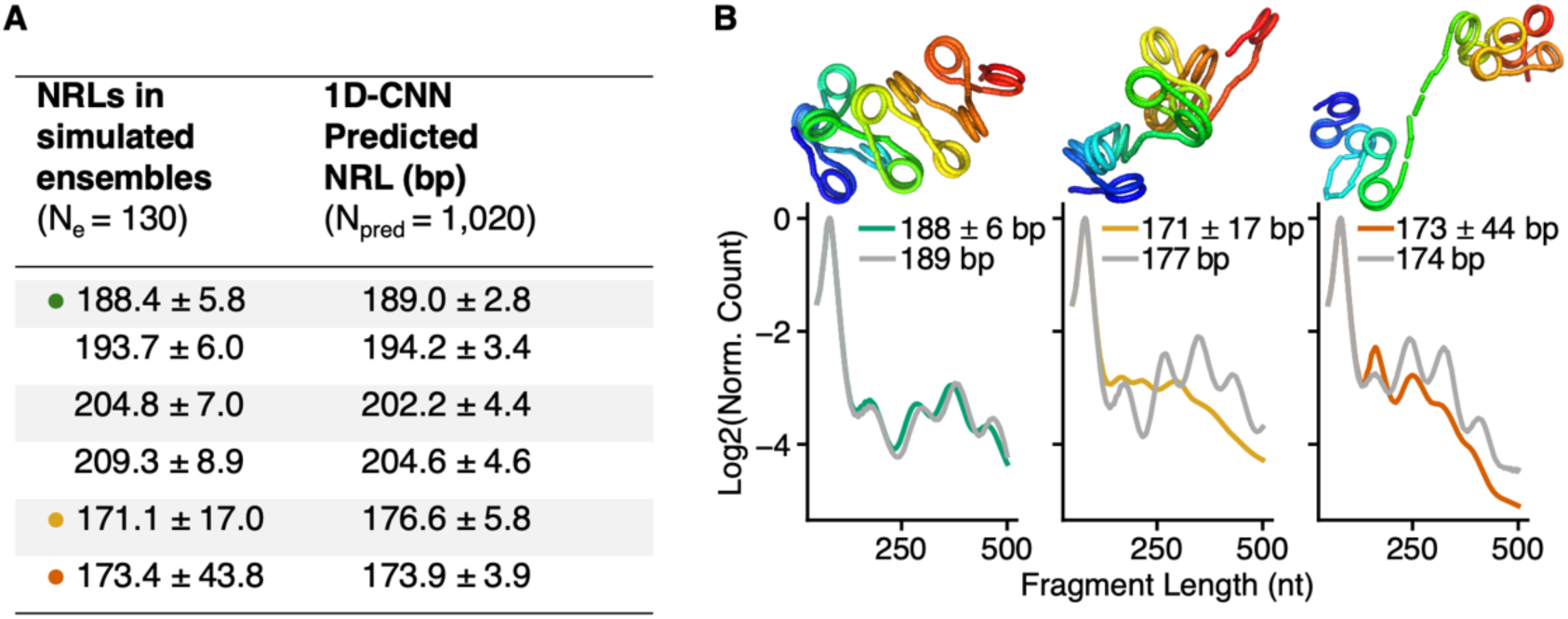
1D-CNN predicts nucleosome spacing in fibers with linker length heterogeneity. (**A**) Numbers are in the form μ±σ where μ is the average and σ is one standard deviation. Ne represents 10 trials of 13 linker DNA segments, Npred represents 102 meso-wlc snapshots for each of 10 trials. (**B**) Gray: the mean FLD from the ensemble with the (homogeneous) NRL predicted by the CNN. Green, yellow, and orange FLDs are the synthetic FLDs calculated from heterogeneous meso-wlc ensembles with low, medium, and high linker length heterogeneity. These FLDs are input into the model trained only on homogeneous linker lengths (Model 3) to derive the predicted NRLs.

### 1D-CNN predicts nucleosome repeat lengths from RICC-seq fragment length distributions consistent with independent experimental data

Finally, we tested the performance of our 1D-CNN trained on ensembles with various NRLs (Model 3), on two real FLDs from experimental RICC-seq data. Each FLD is from an independent biological replicate. For each replicate the fragment length distribution was subset to fragments overlapping with open chromatin (according to ATAC-seq data). We considered only the RICC-seq data that overlapped with ATAC-seq accessibility peaks because we can compute nucleosome spacing from ATAC-seq data in the same cell line (Supplementary Figure S8). Experimental RICC-seq FLDs have weaker higher-order peaks beyond ∼200 nt than predicted because length bias in library preparation and short-read sequencers suppress the efficiency of detecting longer DNA fragments. Higher-order peaks in RICC-seq FLDs are also relatively weak in the experimental data from aggregated ATAC-seq peaks because they are drawn from more open chromatin and because ATAC-seq peaks only cover a small fraction of the genome, leading to a more noisy FLD curve (Figure 7A). The nucleosome repeat length in BJ fibroblasts measured using ATAC-seq is 185.5 bp (Supplementary Figure S8). The CNN predicted an NRL of 182 bp in Replicate 1 (Figure 7A), 3.5 bp from the best-estimate ground truth, and 186 bp in Replicate 2, 0.5 bp from the ground truth from ATAC-seq (Figure 7B). The predicted FLD for Replicate 1 (Figure 7A) has third and fourth peak locations that do not match well with the experimental data, perhaps indicating that the 1D-CNN is not simply predicting the NRL with the most comparable FLD peak locations, but is also learning the unintuitive relationship between chromatin fiber geometric properties and experimental FLDs. The predicted FLD for Replicate 2, however, has peaks in similar locations to the experimental data. This difference between replicates is likely due to the noise limitations of the ATAC-seq FLD curves. This same 1D-CNN also successfully predicted nucleosome spacing from radiation gels (Supplementary Figure S9) (89).

**Figure 7:**
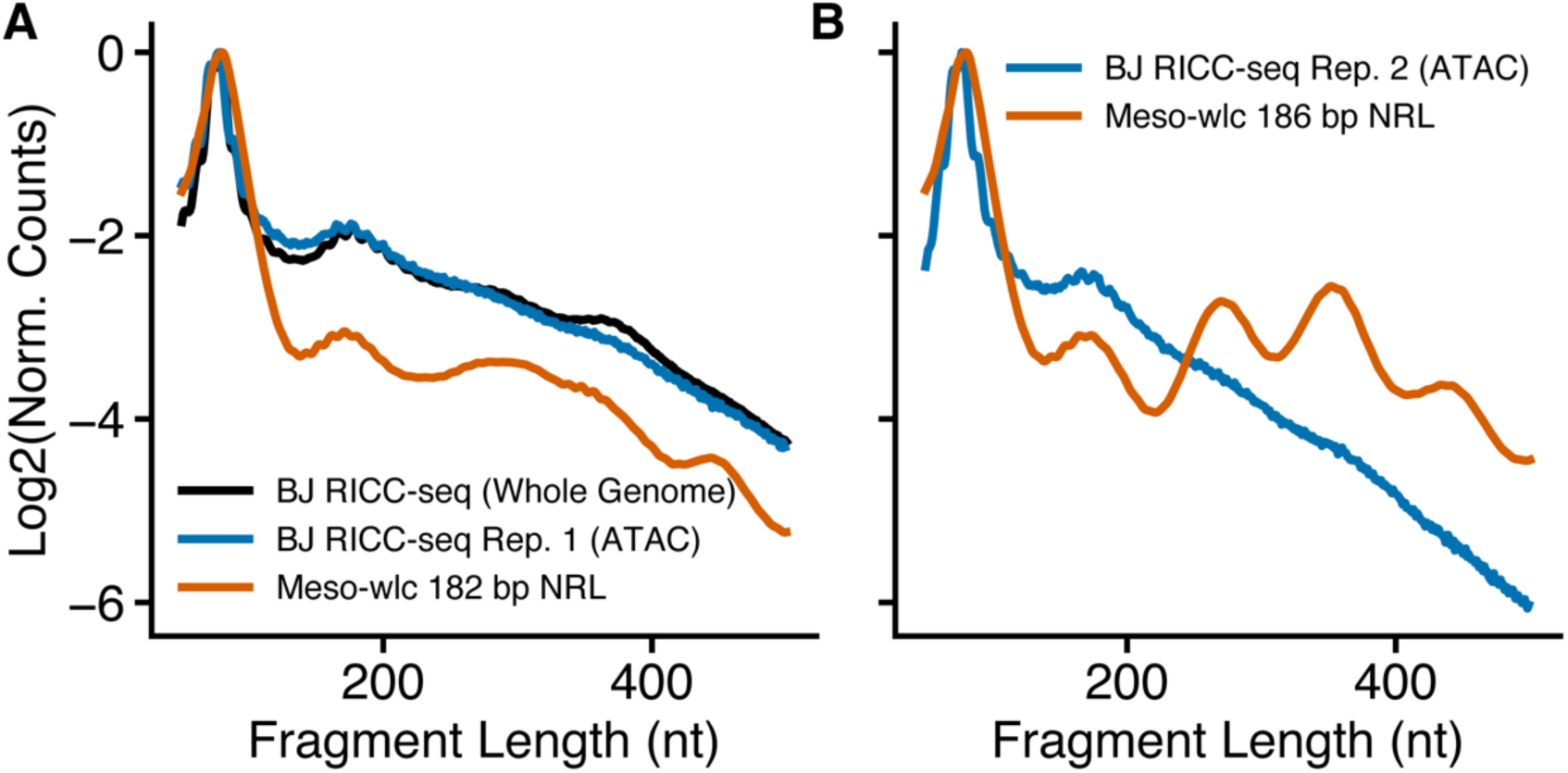
1D-CNN predicts nucleosomal spacing from experimental RICC-seq-derived DNA fragment length distributions. The CNN trained only on simulated data (Model 3) was applied to the RICC-seq data (previously published) (*51*). We analyzed the RICC-seq signal that overlapped with ATAC-seq regions for two biological replicates. (**A**) An NRL of 182 bp was predicted by the CNN for the first biological replicate. The RICC-seq data (blue) is shown alongside the predicted FLD for a meso-wlc ensemble with 182 bp NRL. Whole genome RICC-seq data is shown in black. (**B**) An NRL of 186 bp was predicted by the CNN for the second biological replicate. The RICC-seq data (blue) is shown alongside the predicted FLD for a meso-wlc ensemble with 186 bp NRL. Both predictions are close to 186 bp, the NRL calculated from ATAC-seq data in BJ data (*51*) See Supplementary Figure S8 for calculation of BJ NRL from ATAC-seq data.

## DISCUSSION

We introduced a simple nucleosome geometry- and worm-like chain polymer-based chromatin model that implements coarse-grained DNA elastic parameters including DNA twist, steric interactions, and inter-nucleosome attractive potentials. Using this model, we showed that we can infer key structural features of oligonucleosomes from RICC-seq experiments (51), significantly increasing the impact and usefulness of this method in epigenomic profiling. FLDs contain information about the characteristic distances spanned by DNA self-contacts in folded chromatin, but are insufficient to fully constrain a unique three-dimensional conformation of DNA. Our results demonstrate that despite not being able to fully specify a folded DNA structure, RICC-seq FLDs are nevertheless rich in information about the average structure of the chromatin fibers and the nucleosomes that comprise it. We show that this information can be extracted using our simulation and machine learning platform.

1D-CNNs trained on simulated chromatin structure ensembles performed well at inferring nucleosomal DNA wrapping, NRL, and inter-nucleosome potential from both synthetic and experimental data sets. We successfully predicted NRLs, within six base pairs of ground truth-values, in oligonucleosomes with heterogeneous linker lengths using a model trained only on simulated data with uniform linker lengths (Model 3) (Figure 6). This same model accurately predicts NRLs, within four base pairs of ground-truth values, from real human fibroblast RICC-seq data (Figure 7) (51). We can also simultaneously infer linker length and nucleosome wrapping from synthetic data sets (Model 4), suggesting that the RICC-seq FLD curves are an informative enough data set to allow extraction and deconvolution of both of these structural features.

Whereas several methods besides RICC-seq can measure nucleosome spacing in cell samples (85, 87, 88, 90–92), measuring nucleosomal DNA wrapping *in situ* is more difficult, requiring either very well positioned nucleosomes, which are rare in mammalian cells (92, 93), or single-molecule nucleosome footprinting (85, 87, 88), which currently is still challenging to do at high coverage. RICC-seq has the potential to efficiently measure nucleosome wrapping without requiring single-molecule measurements because the second FLD peak provides information about the average extent of nucleosome wrapping or breathing. Measuring local inter-nucleosome interactions in a sequence-defined manner is also difficult. For example, microscopy (33, 41) does not provide sequence information, while Micro-C and related methods can be prone to artifacts from enzymatic digestion or crosslinking bias (48). We find that the effects of NRL, nucleosomal DNA wrapping, and inter-nucleosome potential strength on chromatin structure (10, 11, 13–15) are all discernible from RICC-seq data when analyzed with the simulation and machine learning approach described here. In combination with meso-wlc, RICC-seq thus becomes a promising technique for measuring these chromatin structure features using a non-enzymatic cleavage mechanism that is orthogonal to other methods, expanding the available toolkit for *in situ* epigenomics.

We designed meso-wlc to be relatively simple, primarily focusing on modeling the geometric features of chromatin and using effective potentials for nucleosome-nucleosome interactions and a coarse-grained worm-like chain model for DNA elasticity. This simple approach is limiting in that meso-wlc cannot be used to explicitly model the effects of molecular components, such as linker histones and particular histone tail modifications. However, this simplicity makes it computationally tractable to make fine-grained adjustments to the NRL and nucleosomal DNA wrapping parameters over a large, but not exhaustive, range of values. Upon introduction of heterogeneity in linker lengths, the possible parameter space becomes very large and cannot be deeply sampled, but it is promising that a neural net trained on homogeneous DNA linkers can still make inferences about the average nucleosome spacing in heterogeneous-linker fibers (Figure 6).

Although we focus on RICC-seq here, the meso-wlc model could also be combined with a model for predicting DNA-DNA contacts measured by other assays. In future applications, meso-wlc could be used to predict signal from other mesoscale chromatin structure probing methods, including Micro-C (46) and Hi-CO (50). Because these methods are assays in which DNA contacts are crosslinked and ligated, DNA-DNA contact prediction models specific to these assays could be defined based on estimates of a physical distance-to-ligated-contact relationship. Furthermore, the neural net-based approach for identifying key features of RICC-seq fragment size distributions could also be applied to ensembles of chromatin structures generated by other simulation platforms—as long as the training set sufficiently samples the parameters of interest.

Overall, pairing mesoscale structural data from sequencing-based assays with meso-wlc simulations boosts the interpretive power of such assays—facilitating the characterization of mesoscale chromatin folding in enough detail to open up new testable hypotheses for understanding the regulation of DNA-based processes by chromatin. These tools provide a new way to gain a more detailed understanding of how the three-dimensional chromatin landscape encountered by chromatin-interacting proteins and the physical behavior of the chromatin fiber are shaped by nucleosome positioning, stability, and modifications.

## DATA AVAILABILITY

The simulation data, intermediate data, and the version of the github code repositories underlying this article are available in Zenodo at https://zenodo.org/doi/10.5281/zenodo.12735673. Scripts are available at https://github.com/riscalab/meso_wlc_manuscript_materials.git. The meso-wlc github repository is available at https://github.com/riscalab/wlcsim.git.

## SUPPLEMENTARY DATA

Supplementary Data are available at NAR Online.

## ACKNOWLEDGEMENTS

We would like to thank Bruno Beltran and Quinn MacPherson for their guidance in adopting kinked tWLC. We would also like to thank Jason Banfelder, Doron Haviv, Ariana Tse, and Joseph Wakim for valuable discussions. We thank the Rockefeller University High Performance Computing Resource Center for computational resources and assistance.

