## Supplemental Information for "Determining mesoscale chromatin structure parameters from spatially correlated cleavage data using a coarse-grained oligonucleosome model"

#### Model Description

##### Steric collision detection

Steric clashes between nucleosomes are detected using the Gilbert-Johnson-Keerthi (GJK) algorithm, which is commonly used for computationally efficient collision detection of convex polyhedra (1; 2). Only collisions within a critical radius were considered where the critical radius is defined as $\sqrt{R_{\text{nuc}}^2 + H_{\text{nuc}}^2}$ (parameters are defined in Supplementary Table S1). Steric collisions were rejected.

##### Inter-nucleosome potential

Nucleosomes favorably interact via contacts mediated by the histone tails' basic residues (3; 4; 5). Cryo-EM images of mono-nucleosomes *in vitro* have shown there are three modes of inter-nucleosome interactions: face-face, face-side, side-side (6). Previous work has investigated the free energies of nucleosome-nucleosome interactions in these three principal modes at 300 K and 150 mM monovalent salt (7). From the minimum inter-nucleosome energies and their respective distances that were previously determined for each interaction orientation (7), we fit a linear fall-off term for that orientation's energy to define a shape of the inter-nucleosome potential in meso-wlc (Supplementary Figure S1).

To weigh the potentials defined above according to the relative orientations of the nucleosomes of interest, we define three coefficients below which capture the extent to which each interaction is face-face, face-side, or side-side.

$$\begin{aligned} \text{Face-face} : k_{\text{ff}} &= |\cos\theta_i||\cos\theta_j| \\ \text{Face-side} : k_{\text{fs}} &= |1-\cos\psi|\mathbf{\Lambda}(|\cos\phi_i|+|\cos\phi_j|) \\ \text{Side-side} : k_{\text{ss}} &= (1-|\cos\phi_i|)(1-|\cos\phi_j|) \end{aligned} \quad \text{(Equation S1)}$$

where

$$\mathbf{\Lambda}(x) = \begin{cases} 2-x & \text{if } x > 1 \\ x & \text{otherwise} \end{cases} \quad \text{(Equation S2)}$$

The orientation angles ($\theta$,$\psi$, $\phi$) are illustrated in Supplementary Figure S2. An interactive gui is available that computes these coefficients for sample nucleosome orientations: `https://github.com/riscalab/meso_wlc_manuscript_materials/tree/main/internucleosome_demo`.

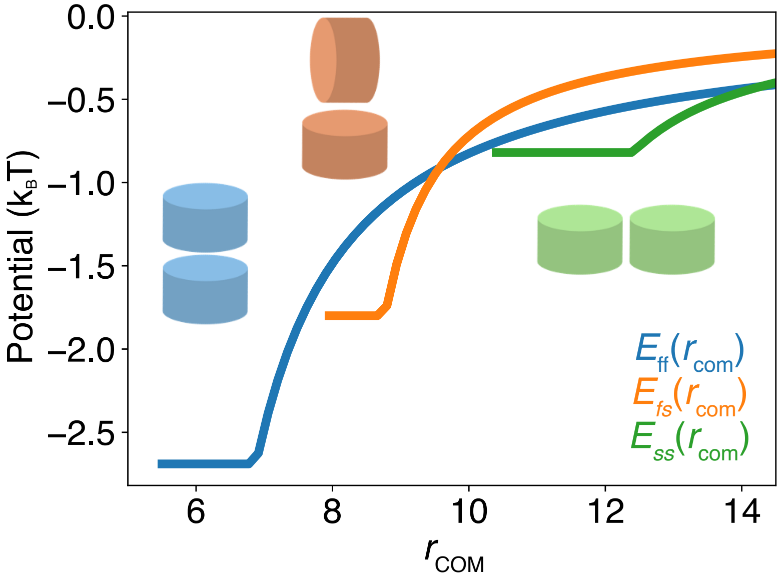


**Supplementary Figure S1: Inter-nucleosome potential is modeled as a function of inter-nucleosome distance.** Potentials are computed for each of the three different inter-nucleosome orientations: face-face (blue), face-side (orange), and side-side (green). The potentials are computed using a linear fall-off term. The minima of the energies at specified distances were previously defined (7).

### Model validation

#### Wrapping parameter selection

Mono-nucleosome structures have been resolved *in vitro* using Cryo-EM (8; 9). One study measured an angle between each linker DNA and the dyad axis in the plane parallel to the nucleosomal disc plane and reported the distribution of these $\alpha$ angles (8) (See Supplementary Figure S3A). A second study resolved mono-nucleosome structures from Xenopus egg extract (9). They compute a "face angle" that is equivalent to -2$\alpha$ (See Supplementary Figure S3A). These face angles are computed for each resolved 3D structure class along with the percent of particles that belong to each class, enabling the computation of a weighted average face angle per experimental condition. Two of these experimental conditions are considered herein: 1.) "Freed" interphase mono-nucleosomes that are MNase digested before crosslinking and 2.) Interphase mono-nucleosomes that are MNase digested after crosslinking (9). It is possible that cross-linking before digestion (experimental condition 2) may result in more crosslinking between the DNA and nucleosome core complex due to interactions within the pre-digested complex. This would correspond to higher $\alpha$ values. By contrast, MNase digestion before crosslinking (experimental condition 1) may result in over-digested mono-nucleosomes which would have lower $\alpha$ values. We use the $\alpha$ values directly or indirectly reported in these two studies to fit a relevant default nucleosomal DNA wrapping parameter for meso-wlc. We carried out simulations with various wrapping parameter values and determined that a wrapping parameter of 127 bp reasonably aligns with the Cryo-EM results. Results for four simulated experimental conditions (122, 127, 132, and 147 bp wrapped around the nucleosome core) are provided in the main text, but additional simulations are included in Supplementary Figure S4.

Validating this parameter selection against sedimentation assay data further supports that 127 bp is a reasonable nucleosomal DNA wrapping parameter. We validated against sedimentation assays where the histone tails were removed (10). This eliminated the need to calibrate both wrapping and inter-nucleosome potential simultaneously. Without histone tails, the effective inter-nucleosome potential is $0 k_B T$. Dorigo et al. ran sedimentation assays that determined that fibers with 12 nucleosomes and an NRL of 177 bp have sedimentation values of 39.6 Svedberg and 42.6 Svedberg in 0.6 mM $Mg^{2+}$ and 1.0 mM $Mg^{2+}$ *in vitro*, respectively. Exact values were extracted using WebPlotDigitizer (v4.8) (11). These ionic conditions are consistent with the physiological range (12). We ran simulations of 12mers with the same NRL, no inter-nucleosome potential, and a range of wrapping parameters. We computed the predicted sedimentation coefficients using Equation S11 and found a wrapping parameter of 127 bp to be consistent with the experimental sedimentation coefficients (Supplementary Figure S5).

These rigorous comparisons with experimental data support the selection of 127 bp as the default nucleosomal

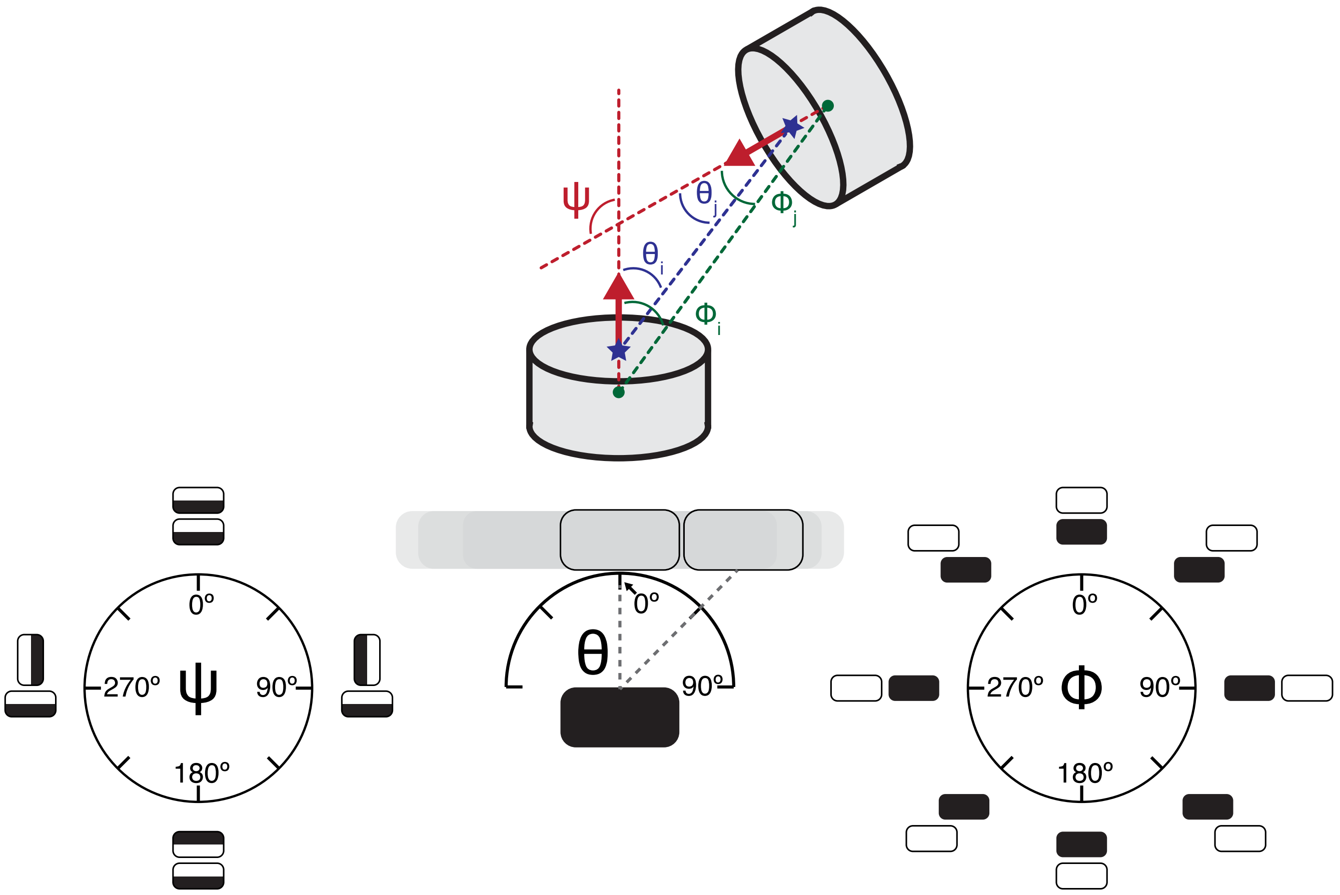


**Supplementary Figure S2: Inter-nucleosome angles are defined.** (Top) The rotational angle, $\psi$, describes the torsional angle between two nucleosome face normal vectors (depicted as red arrows). The angle of inclination, $\theta_x$, is the angle between nucleosome $x$'s face normal and the distance vector between a pair of nucleosomes' nearest face centers. The center of the nearest faces are depicted as blue stars. The angle between nucleosomes $x$'s face normal vector and the distance vector connecting the two centers of mass is $\phi_x$. The center of mass of each nucleosome is depicted as a green circle. (Bottom) The $\psi$ chart shows all possible rotational angles. The $\psi$ angle distinguishes between directions of face-to-face stacking. The "head" and "tail" sides of each nucleosome are distinguished by shading. Head-to-tail stacking is described by $\psi = 0°$, head-to-head stacking is described by $\psi = 180°$. The $\psi$ angle is used to define the face-side coefficient. The $\theta$ chart shows a range of possible angles. Nucleosome $x$ is depicted in black and the other nucleosomes are unshaded. Two reference nucleosomes are shown with $\psi = 0°$ and the shaded area depicts other possible nucleosome positions with the same $\psi$ angle. The $\theta$ angle is used to define the face-face coefficient. For perfect face-face stacking, $\theta = 0°$, and $\theta$ approaches $90°$ as the inclination between nucleosomes increases. The $\phi$ chart shows all possible angles. Here, nucleosome $x$ is depicted in black and the reference nucleosome is unshaded. All nucleosome pairs are depicted with a fixed $\psi$, $\psi = 0°$. The $\phi$ angle is used to define both the side-side and the face-side coefficient. These inter-nucleosome angles, $\psi$, $\phi$, and $\theta$, are collectively used to compute the orientation coefficients defined in Equation S1.

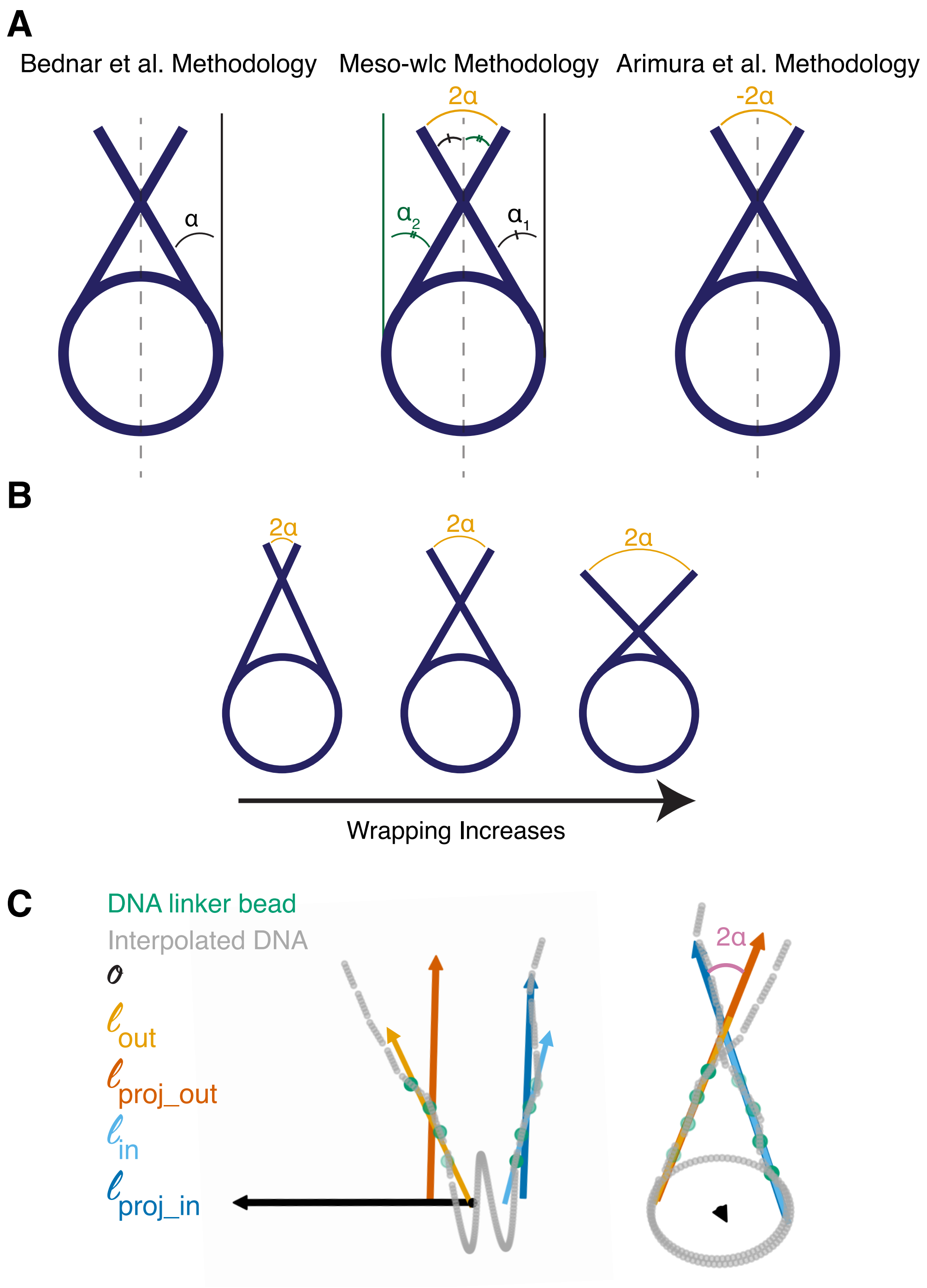


**Supplementary Figure S3: Calculation of the entry-exit angle.** (**A**) The entry-exit angle, $\alpha$ as defined in (8). Here $\alpha$ is the angle between each linker DNA and the dyad axis in the plane parallel to the nucleosomal disc plane. Note that $\alpha_1$ does not necessarily equal $\alpha_2$, these numbers are averaged to compute $\alpha$. (**B**) The entry-exit angle increases as the amount of DNA wrapped around the nucleosome core increases. (**C**) The orthonormal vector, $\vec{o}$ is depicted in black. It points through the face of the nucleosome core particle in the direction from DNA entry to DNA exit. The first principal components of the entry and exit linker DNA segments are depicted in light blue ($\vec{l}_{in}$) and gold ($\vec{l}_{out}$), respectively. These vectors are then projected onto the plane perpendicular to $\vec{o}$. These projected vectors are shown in dark blue ($\vec{l}_{\text{proj\_in}}$) and orange ($\vec{l}_{\text{proj\_out}}$) and the angle between these two vectors is computed, $2\alpha$.

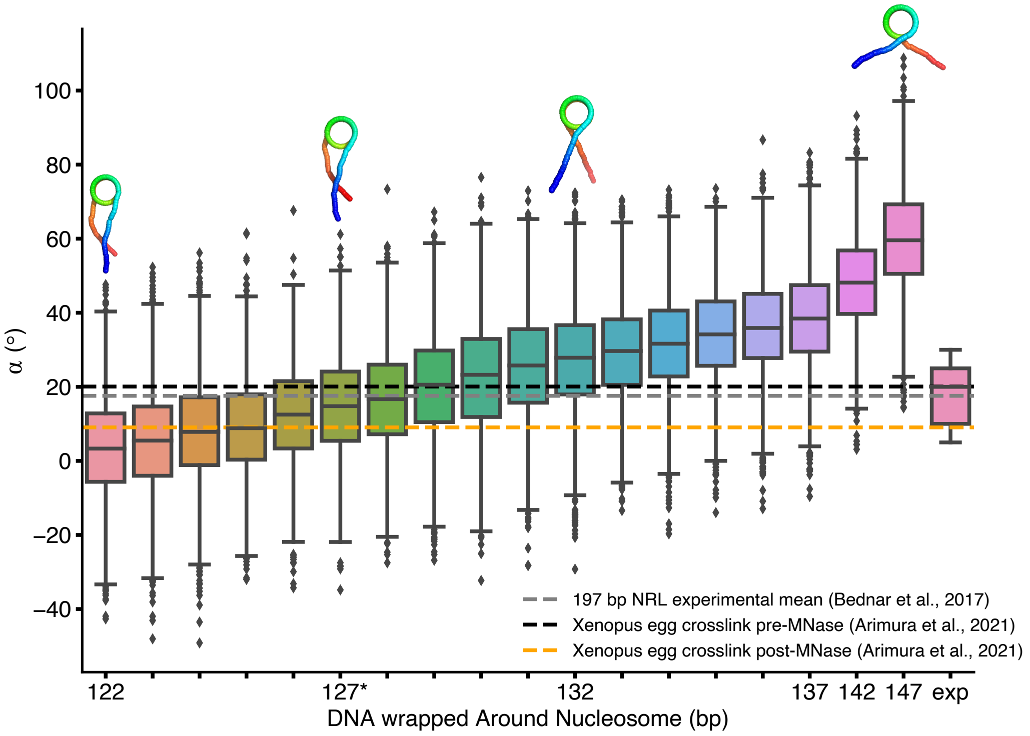


**Supplementary Figure S4: A nucleosome wrapping parameter of 127 bp yields consistent entry-exit angles ($\alpha$) as Cryo-EM data of mono-nucleosomes.** The asterisk denotes the default parameter selected for subsequent simulations. Entry-exit angles calculated from simulated ensembles of mononucleosomes are compared against Cryo-EM data (8; 9). The distribution of entry-exit angles labeled 'exp' is from Bednar et al., 2017 (8).

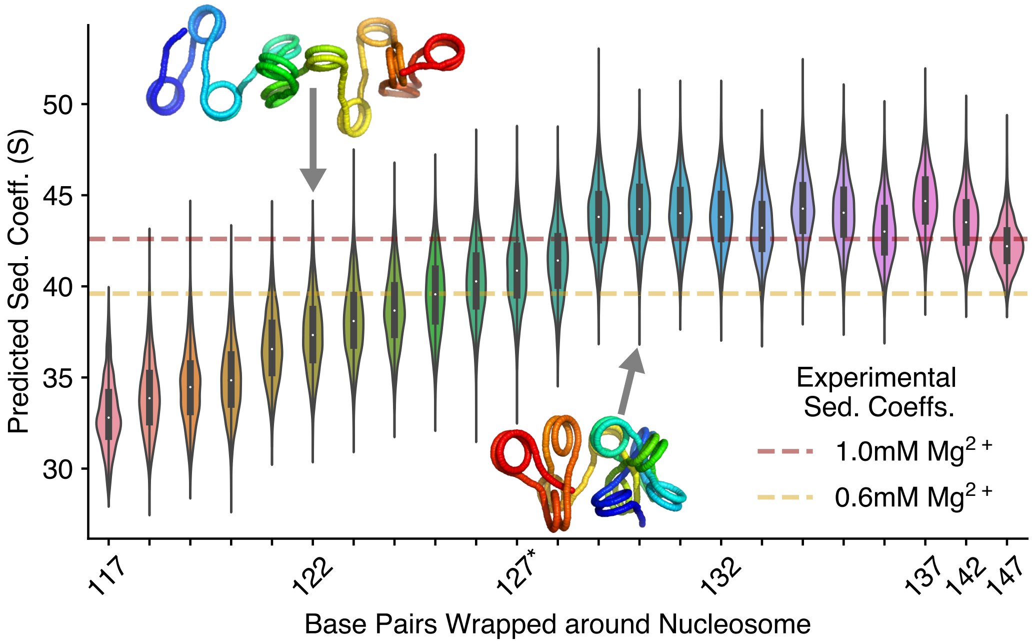


**Supplementary Figure S5: A wrapping parameter selection of 127 bp yields a predicted sedimentation coefficient that is consistent with experimental data** (10). The asterisk denotes the default parameter selected for subsequent simulations. The experimental data comes from mutant nucleosome arrays without histone tails. Sedimentation coefficients for assays at 0.6 mM and 1.0 mM magnesium are included because these concentrations are consistent with physiological salt (12).

DNA wrapping parameter. The user may select a different nucleosomal DNA wrapping parameter.

#### Equilibration and exploration of conformational space

When running simulations, it can be difficult to assess whether the system has been run to equilibrium or is stuck in a long-living metastable state (13; 14). Here, we include an analysis of a representative simulation with a 187 bp NRL. We evaluate how well the conformational space is explored. First, we performed multiple trials of each simulation to decrease the chances of evaluating an unrepresentative meta-stable state. Fifteen trials were performed. Autocorrelation analysis also offers a way to quantify the effective number of uncorrelated samples in a dataset (15). We evaluate the autocorrelation of the distance between the center of the first and last nucleosomes. We do this by computing the Pearson correlation coefficient of these distances at time t and t+x where x is the lag interval (in units of snapshots). We then fit an exponential of the form $ae^{-x/\tau} + b$ to the plot of the Pearson correlation coefficients vs. the lag interval. We find that after an initial burn in period of 10 snapshots, the $\tau = 0.363$. This indicates that each of the 291 snapshots per trial (4,365 across the 15 trials) are sufficiently independent. (Supplementary Figure S6A).

Examining the worm-like chain energies for one of the fifteen trials shows that, after a burn in period of 10 snapshots (red), the fiber is equilibrated (Supplementary Figure S6B). For all simulations considered herein, at least the first 10 snapshots were excluded from analysis. After the first 10 snapshots, the fiber explores the conformational space, as evidenced by changes in the predicted sedimentation coefficient (Supplementary Figure S6C). If the fiber was stuck in a meta-stable state, the predicted sedimentation coefficient would remain constant.

To asses sampling, we can observe the sampling efficiency by looking at the distribution of inter-nucleosome contacts distances (16). By looking at the distances between 1.) The 5th and 7th nucleosomes vs. the 6th and 8th nucleosomes and 2.) The 1st and 11th vs. the 2nd and 12th nucleosomes, we observe that the simulated structures sample the conformational space well and the simulations are not getting stuck in local minima (Supplementary Figure S6D).

### Model energy

The potential energy of the fiber is given by:

$$E(\{\vec{r}, \vec{u}, \vec{w}\}) = \sum_{i=2}^{N} \left[ \frac{\varepsilon_b(\delta)}{2\Delta} |\vec{u}_i - \vec{u}_{i-1} - \eta(\delta)\vec{R}_i^{\perp}|^2 + \frac{\varepsilon_{\parallel}(\delta)}{2\Delta} |\vec{R}_i \cdot \vec{u}_{i-1} - \Delta\gamma(\delta)|^2 + \frac{\varepsilon_{\perp}(\delta)}{2\Delta} |\vec{R}_i^{\perp}|^2 + \frac{\varepsilon_t(\delta)}{2\Delta}\omega_t^2 \right] + E_{\text{sterics}} + E_{\text{internuc}} \quad \text{(Equation S3)}$$

where

$$\vec{R}_i = \vec{r}_i - \vec{r}_{i-1} \quad \text{(Equation S4)}$$

$$\vec{R}_i^{\perp} = \vec{R}_i - (\vec{R}_i \cdot \vec{u}_{i-1})\vec{u}_{i-1} \quad \text{(Equation S5)}$$

$$\omega_t = \arctan\left(\frac{\vec{v}_{i-1} \cdot \vec{w}_i - \vec{w}_{i-1} \cdot \vec{v}_i}{\vec{w}_{i-1} \cdot \vec{w}_i + \vec{v}_{i-1} \cdot \vec{v}_i}\right) \quad \text{(Equation S6)}$$

$$\vec{w}_i = \vec{v}_i \times \vec{u}_i, \forall i \in [1, N] \quad \text{(Equation S7)}$$

Here, $\varepsilon_b$ is the bending modulus, $\eta$ is the bend-shear coupling parameter, $\varepsilon_{\parallel}$ is the stretching modulus, $\gamma$ is the ground-state segment length between consecutive beads (where $\vec{R}_i$ is the actual segment length between consecutive beads), $\varepsilon_{\perp}$ is the shearing modulus, and $\varepsilon_t$ is the twisting modulus. The material moduli ($\gamma$, $\eta$, $\varepsilon_{\parallel}$, $\varepsilon_b$, $\varepsilon_{\perp}$, $\varepsilon_t$) are defined as a function of the user-defined discretization, $\delta$. The moduli are also scaled by $\Delta$, the distance at equilibrium between beads. The process of computing the moduli for a given $\delta$ has been previously determined (17; 18; 19). In this work, the $\delta$ is approximately 8 base pairs per bead. The discretization used for all simulations is shown in Equation S8. In Equation S3, $E_{\text{sterics}}$ is only nonzero if the fiber is initialized with a steric clash. $E_{\text{sterics}} = 3k_BT * N_{\text{clash}}$, where $N_{\text{clash}}$ represents the number of steric clashes. MC moves that introduce a steric

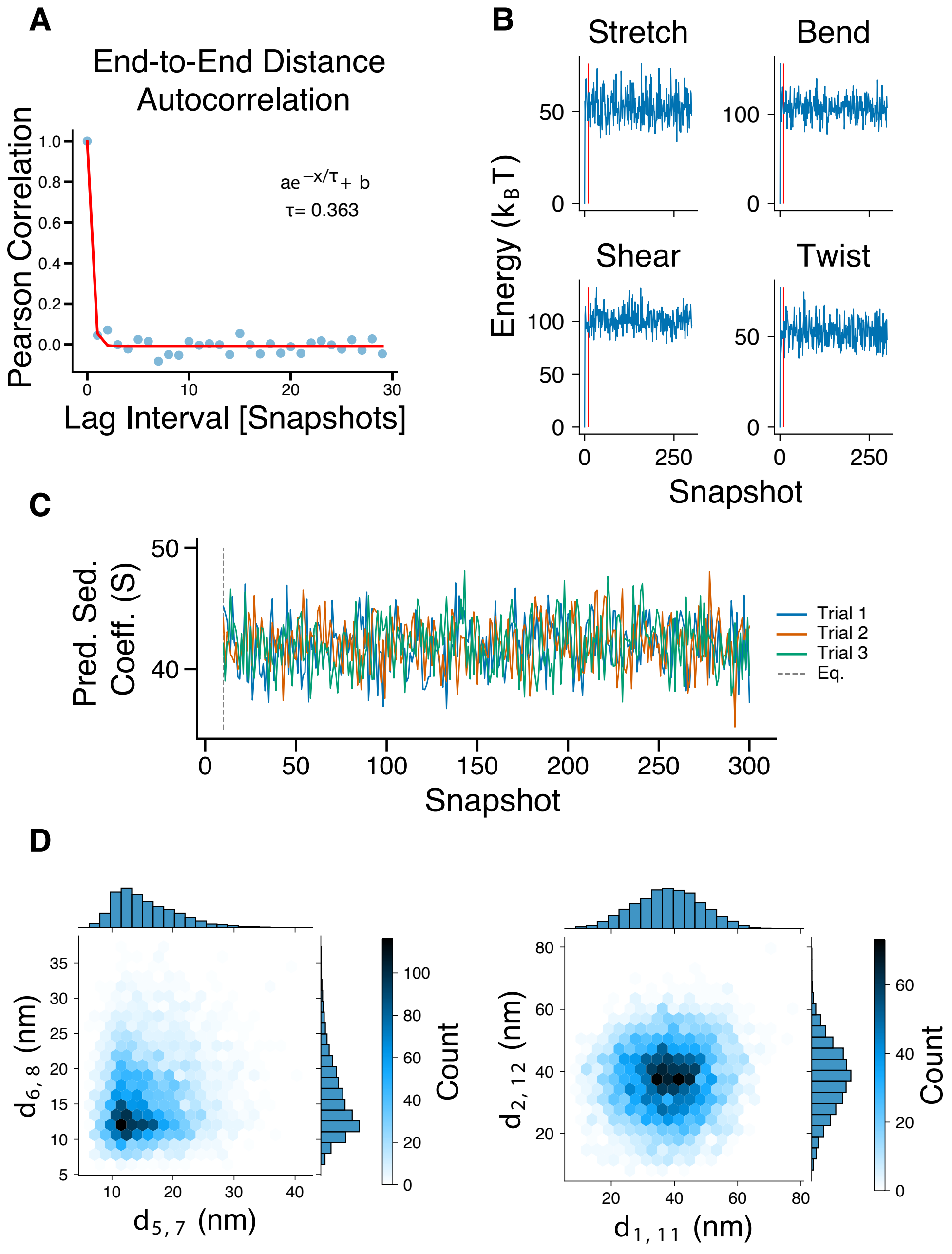


**Supplementary Figure S6: Meso-wlc simulations yield well-sampled ensembles with end-to-end distances that are not autocorrelated.** Simulations shown here are from a fiber with an NRL of 187 bp. 301 snapshots are collected per trial and the first 10 are discarded to allow for a burn-in time. (**A**) End-to-end distances between snapshots are not autocorrelated. Results are averaged over 15 trials. (**B**) One trial is shown. Fiber continues sampling after a burn in period of 10 snapshots. Red vertical line depicts 10 snapshots. (**C**) The predicted sedimentation coefficient data of three trials shows that trials do not get stuck in local energy minima. (**D**) Inter-nucleosome distance analysis shows that conformational space is well-sampled. (Left) N-N+2 distances are shown between the 5th and 7th nucleosomes and the 6th and 8th nucleosomes. (Right) N-N+10 distances are shown between the 1st and 11th and 2nd and 12th nucleosomes. In both panels, the unimodal distributions are consistent with efficient sampling whereas multi-modal distributions would be consistent with the different trials converging to two or more meta-stable states.

clash are rejected before the potential energy of the fiber is computed.

$$N_{\text{beads per linker}} = \text{round}(N_{\text{base pair per linker}}/8) \qquad \text{(Equation S8)}$$

For example, for a linker length of 68bp, there would be 9 beads per linker (round(68/8)=round(8.5)), and the resulting $\delta$ would be 7.56 base pairs per bead. The corresponding $\Delta$ would be 7.56*$z$ nm (where $z$ is the length per base pair as reported in Supplementary Table S1), or 2.51 nm. When a non-integer discretization value ($\delta$) is used, the appropriate material moduli are interpolated from material moduli previously calculated for integer $\delta$ values (18).

The inter-nucleosome potential is described in the section above, but is defined mathematically as:

$$E_{\text{internuc}}(\vec{r}_{com_i}, \vec{r}_{com_j}) = \sum_{j>1}^{N_{\text{nuc}}} \sum_{i=1}^{N_{\text{nuc}}} k_{\text{ff}} * E_{\text{ff}}(r_{\text{com}_{ij}}) + k_{\text{fs}} * E_{\text{fs}}(r_{\text{com}_{ij}}) + k_{\text{ss}} * E_{\text{ss}}(r_{\text{com}_{ij}}) \qquad \text{(Equation S9)}$$

$$r_{com_{ij}} = \sqrt{||\vec{r}_{com_i} - \vec{r}_{com_j}||} \qquad \text{(Equation S10)}$$

Here $r_{\text{com}}$ is the distance between the center of masses of nucleosome $i$ and nucleosome $j$. The number of nucleosomes in the simulated fiber is denoted as $N_{\text{nuc}}$. The coefficients $k_{\text{ff}}$, $k_{\text{fs}}$, $k_{\text{ss}}$ are given by Equation S1. The orientation-specific inter-nucleosome potentials ($E_{\text{ff}}(r)$, $E_{\text{fs}}(r)$, $E_{\text{ss}}(r)$) are computed as shown in Supplementary Figure S1.

### Measurements

#### Sedimentation coefficient

Sedimentation coefficients are calculated *in vitro* by measuring the relative centrifugation rates of nucleosomal arrays. In analytical ultracentrifugation, the speed of centrifugation correlates with the compaction of the nucleosomal array; more compact fibers have higher sedimentation coefficients (20). *In silico* sedimentation coefficients are estimated as a function of the pairwise distances between nucleosomes (21; 22; 23). We computed predicted sedimentation coefficients using the Kirkwood-Bloomfield formulation (24; 25):

$$S_{\text{pred}} = S + \frac{R_{\text{c}}S}{N_{\text{nuc}}} \sum_i \sum_j \frac{1}{r_{com_{ij}}} \qquad \text{(Equation S11)}$$

where $S_{\text{pred}}$ is the predicted sedimentation coefficient, $R_{\text{c}}$ is the hydrodynamic radius of the nucleosome core (5.46 nm) (21), $S$ is the sedimentation coefficient for a mono-nucleosome at $20°$ ($S = 11.1$ Svedberg, where 1 Svedberg = $10^{-13}s$) (23; 26; 27).

#### Entry-exit angle

For each nucleosome, The first 4 beads before and after the nucleosome core particle kink are considered. The first principal component of the 3D coordinates, $\vec{l}_{\text{pre\_in}}$, is computed for the 4 beads before the nucleosome core particle. If the resulting vector is not pointing in the direction from the nucleosome core outwards towards the linker, it is multiplied by -1, else it is used as is to define the entry linker vector, $\vec{l}_{\text{in}}$.

$$\vec{l}_{\text{in}} = \begin{cases} \vec{l}_{\text{pre\_in}}, & \text{if } \vec{l}_{\text{pre\_in}} \text{ points in direction from nucleosome core outwards.} \\ -\vec{l}_{\text{pre\_in}}, & \text{otherwise.} \end{cases} \qquad \text{(Equation S12)}$$

The process defined above is also used to compute the exit linker vector $\vec{l}_{\text{out}}$ from the 4 beads after the nucleosome core particle kink. These vectors ($\vec{l}_{\text{in}}$ and $\vec{l}_{\text{out}}$) are then projected onto the plane perpendicular to the nucleosome's orthonormal vector, $\vec{o}$. Note, $\vec{o}$ is defined to point in the direction from DNA entry to DNA exit as shown in Supplementary Figure S3C.

$$\vec{l}_{\text{proj\_in}} = \vec{l}_{\text{in}} - \frac{\vec{l}_{\text{in}} \cdot \vec{o}}{||\vec{o}||^2}\vec{o} \qquad \text{(Equation S13)}$$

$$\vec{l}_{\text{proj\_out}} = \vec{l}_{\text{out}} - \frac{\vec{l}_{\text{out}} \cdot \vec{o}}{||\vec{o}||^2}\vec{o} \qquad \text{(Equation S14)}$$

$$2\alpha = \text{arccos}\left(\frac{\vec{l}_{\text{proj\_in}} \cdot \vec{l}_{\text{proj\_out}}}{|\vec{l}_{\text{proj\_in}}||\vec{l}_{\text{proj\_out}}|}\right) \qquad \text{(Equation S15)}$$

#### BJ fibroblast nucleosome repeat length

The nucleosome repeat length was calculated using eight biological replicates of ATAC-seq data from BJ fibroblasts (28). Previously published RICC-seq and ATAC-seq data (28) are available in the NCBI Gene Expression Omnibus (29) under accession number GSE81807. The fragment length distributions were analyzed for each replicate. The distributions were subset to fragment lengths greater than 140 bp (to filter out sub-nucleosomal fragments) and less than 500 bp. For each replicate, LOWESS smoothing with a parabolic term and a window size of 80 bp was applied to the log10 transformed fragment length distribution. Local maxima were identified corresponding to the mono-nucleosome and di-nucleosome peak. The distance between these two peaks was computed for each replicate and averaged between replicates to yield an NRL of 185.5 bp (See Supplementary Figure S8).

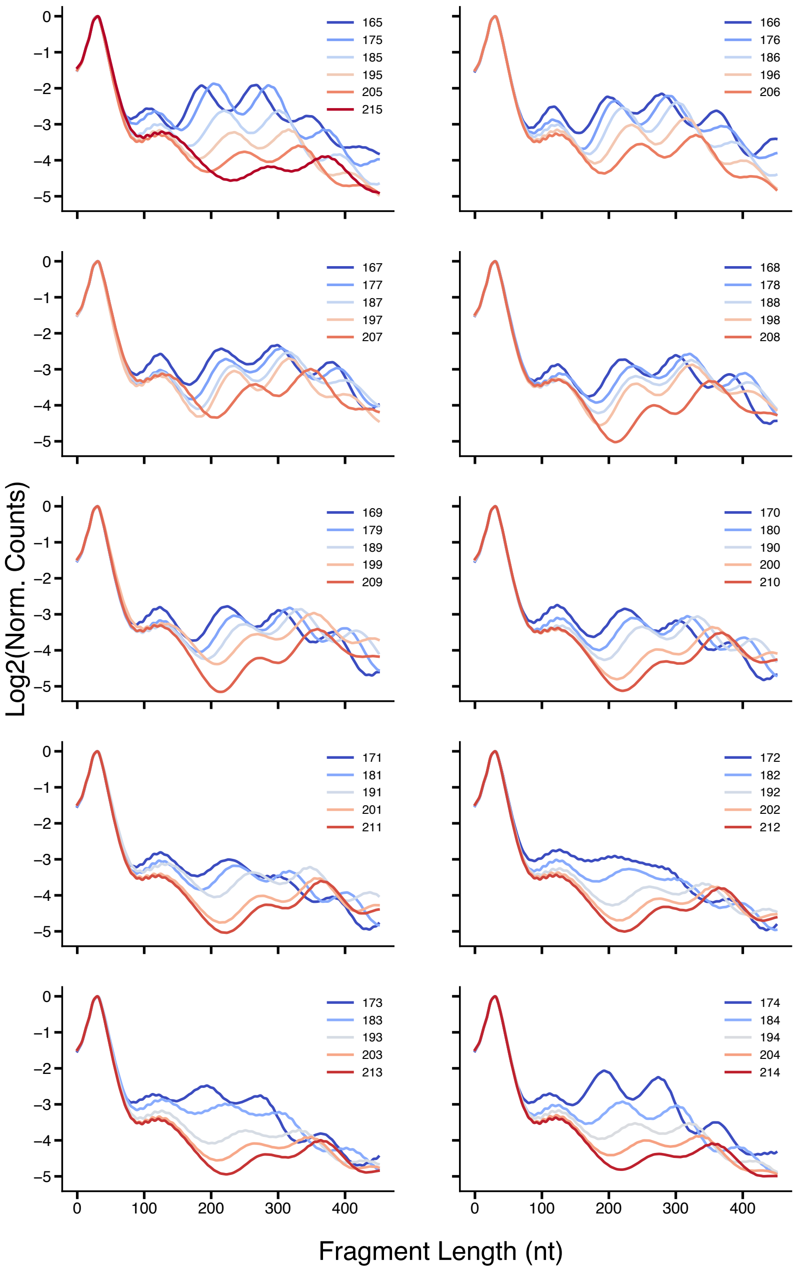


**Supplementary Figure S7: The nucleosome repeat length has a dramatic effect on the predicted fragment length distribution.** The plotted NRL is indicated in the legend. 4,365 snapshots are summarized for each NRL. In general, NRL length is negatively correlated with density in the 3rd-5th predicted RICC peaks. See Supplementary Table S5 for additional simulation details.

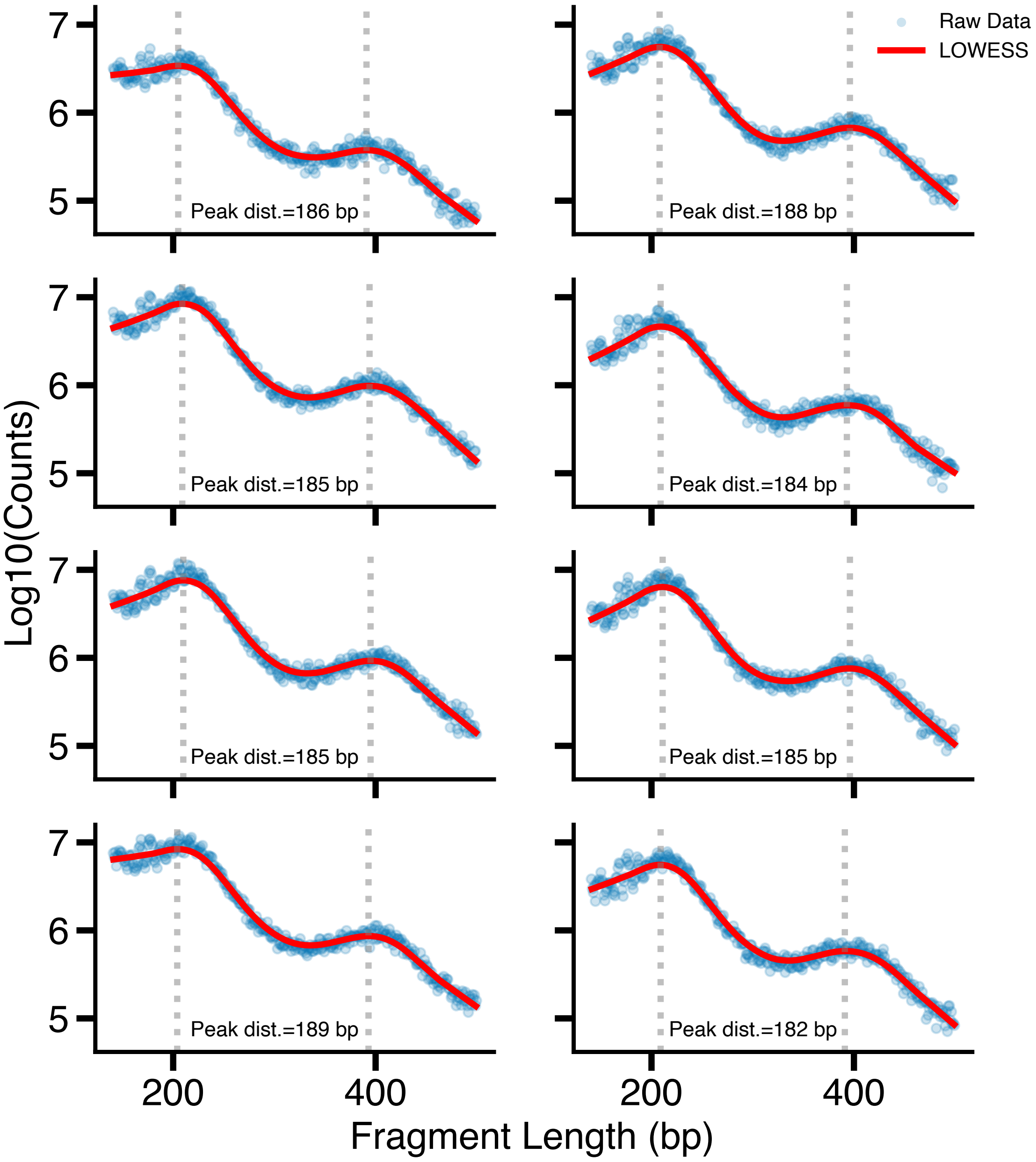


**Supplementary Figure S8: ATAC-seq data show BJ fibroblasts have an NRL of 185.5 bp.** The raw histogram data for each of eight replicates is plotted (blue dots) along with the LOWESS smoothed data (red line). The dotted lines represent local maxima. The first maximum is the mono-nucleosome peak and the second represents the di-nucleosome peak. The distance between the two maxima (the peak distance) is the NRL. The NRL is indicated on each subplot. The average of the eight NRLs shown is 185.5 bp.

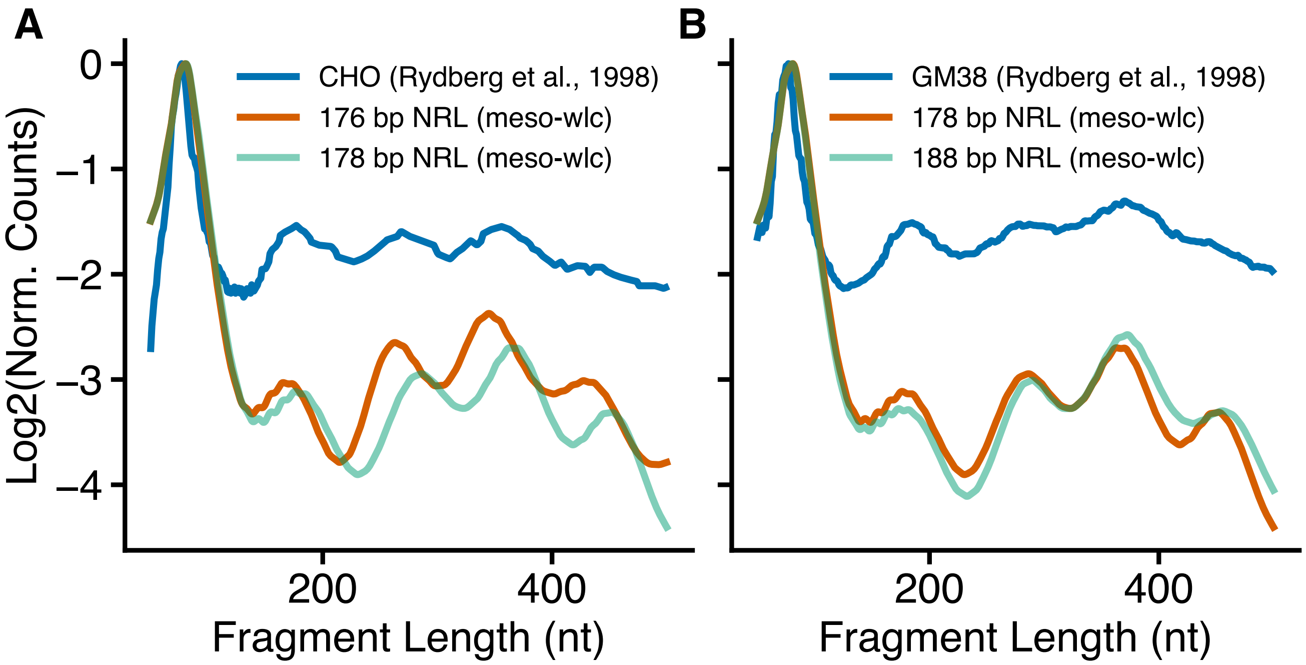


**Supplementary Figure S9: Accurate nucleosome spacing is predicted from radiation gel by 1D-CNN trained on simulated chromatin ensembles.** Data was extracted from Rydberg et al. 1998 (30) using WebPlotDigitizer (v4.8) (11). Experimental data is plotted in blue. FLDs from meso-wlc ensembles with the NRL predicted by the CNN are shown in orange. FLDs from meso-wlc ensembles with the cell line of interest's reported NRL are shown in green. (**A**) Our model predicts an NRL of 176 bp from radiation gel data from mitotic CHO cells. This is within 2 bp of the 178 bp NRL reported for these cells (30; 31). (**B**) Our model predicts an NRL of 178 bp from GM38 human fibroblast data. The FLDs calculated from meso-wlc structure ensembles with NRLs of 178 bp and 188 bp are shown. An NRL of 188 bp has been reported for similar cell types (30; 31).

**Supplementary Table S1:** Coarse-grained and interpolated fiber parameters

| Parameter | Description | Value | Ref. |
|---|---|---|---|
| $R_{\text{nuc}}$ | Nucleosome radius | 5.2 nm | |
| $H_{\text{nuc}}$ | Nucleosome height | 5.5 nm | |
| $R_{\text{c}}$ | Hydrodynamic radius of nucleosome core | 5.46 nm | (21) |
| $S$ | Sedimentation coefficient for a mono-nucleosome at 20°C | 11.1 Svedberg | (23; 26; 27) |
| $z$ | Distance between base pairs | 0.332 nm | (32) |
| $l_P$ | DNA Persistence length | 50 nm | (33; 34) |
| $R_{\text{DNA}}$ | DNA radius | 1 nm | (35; 36) |
| $\omega$ | DNA twist (radians/bp) | $\frac{2\pi}{10.5}$ | (37) |
| $\beta$ | Phase difference between plus and minus strand (radians) | $0.77\pi$ | (38; 39) |

**Supplementary Table S2:** Monte Carlo parameters

| Move Type | Number per MC Step |
|---|---|
| Crank shaft | 30 |
| Pivot | 30 |
| Bead translation | 60 |
| Bead rotation | 60 |
| Chain rotation | 5 |
| Chain translation | 6 |

**Supplementary Table S3:** Simulation parameters: mono-nucleosome simulations used to calibrate nucleosomal DNA wrapping

| Index | Fig. Number | NRL (bp) | Wrapping (bp) | Trials | Steps/ Snapshot | Total Snapshots | Burn-in | Beads/Linker |
|---|---|---|---|---|---|---|---|---|
| 1 | 2A, S4 | 197 | 122 | 15 | 2,093 | 201 | 10 | 8 |
| 2 | S4 | 197 | 123 | 15 | 2,093 | 201 | 10 | 9 |
| 3 | S4 | 197 | 124 | 15 | 2,093 | 201 | 10 | 9 |
| 4 | S4 | 197 | 125 | 15 | 2,093 | 201 | 10 | 9 |
| 5 | S4 | 197 | 126 | 15 | 2,093 | 201 | 10 | 9 |
| 6 | 2A, S4 | 197 | 127 | 15 | 2,093 | 201 | 10 | 9 |
| 7 | S4 | 197 | 128 | 15 | 2,093 | 201 | 10 | 9 |
| 8 | S4 | 197 | 129 | 15 | 2,093 | 201 | 10 | 9 |
| 9 | S4 | 197 | 130 | 15 | 1,872 | 201 | 10 | 8 |
| 10 | S4 | 197 | 131 | 15 | 1,872 | 201 | 10 | 8 |
| 11 | 2A, S4 | 197 | 132 | 15 | 1,872 | 201 | 10 | 9 |
| 12 | S4 | 197 | 133 | 15 | 1,872 | 201 | 10 | 9 |
| 13 | S4 | 197 | 134 | 15 | 1,872 | 201 | 10 | 9 |
| 14 | S4 | 197 | 135 | 15 | 1,872 | 201 | 10 | 9 |
| 15 | S4 | 197 | 136 | 15 | 1,872 | 201 | 10 | 9 |
| 16 | S4 | 197 | 137 | 15 | 1,872 | 201 | 10 | 9 |
| 17 | S4 | 197 | 142 | 15 | 1,652 | 201 | 10 | 9 |
| 18 | 2A, S4 | 197 | 147 | 15 | 1,432 | 201 | 10 | 8 |

* All included simulations are of mono-nucleosomes.

**Supplementary Table S4:** Simulation parameters: dodecamer simulations used to calibrate inter-nucleosome potential

| Index | Fig. Number | NRL (bp) | Inter-nucleosome Potential ($k_BT$) | Trials | Steps/ Snapshot | Total Snapshots | Burn-in | Beads/Linker |
|---|---|---|---|---|---|---|---|---|
| 1 | 2B | 207 | 2.7 | 10 | 11,567 | 201 | 10 | 8 |
| 2 | 2B | 207 | 5.4 | 10 | 11,567 | 201 | 10 | 8 |
| 3 | 2B | 207 | 8.1 | 10 | 11,567 | 201 | 10 | 8 |
| 4 | 2B | 207 | 10.8 | 10 | 11,567 | 201 | 10 | 8 |
| 5 | 2B | 207 | 13.5 | 10 | 11,567 | 201 | 10 | 8 |
| 6 | 2B | 207 | 16.2 | 10 | 11,567 | 201 | 10 | 8 |
| 7 | 2B | 207 | 18.9 | 10 | 11,567 | 201 | 10 | 8 |
| 8 | 2B, 5A-B | 188 | 0 | 3 | 11,567 | 201 | 50 | 8 |
| 9 | 2B, 5B | 188 | 2.7 | 3 | 11,567 | 201 | 50 | 8 |
| 10 | 2B, 5B | 188 | 5.4 | 3 | 11,567 | 201 | 50 | 8 |
| 11 | 2B, 5B | 188 | 8.1 | 3 | 11,567 | 201 | 50 | 8 |
| 12 | 2B, 5A-B | 188 | 10.8 | 3 | 11,567 | 201 | 50 | 8 |
| 13 | 2B, 5B | 188 | 13.5 | 3 | 11,567 | 201 | 50 | 8 |
| 14 | 2B, 5B | 188 | 16.2 | 3 | 11,567 | 201 | 50 | 8 |
| 15 | 2B, 5B | 188 | 18.9 | 3 | 11,567 | 201 | 50 | 8 |
| 16 | 2B, 5A-B | 188 | 21.6 | 3 | 11,567 | 201 | 50 | 8 |

* All included simulations are of dodecamers (12 nucleosomes) with 127 bp nucleosomal DNA wrapping.

**Supplementary Table S5:** Simulation parameters: dodecamer simulations with various nucleosome repeat lengths

| Index | Fig. Number | NRL (bp) | Trials | Steps/ Snapshot | Total Snapshots | Burn-in | Beads/Linker |
|---|---|---|---|---|---|---|---|
| 1 | 2C, 4E, S7 | 165 | 15 | 7,271 | 301 | 10 | 5 |
| 2 | 2C, 4E, S7 | 166 | 15 | 7,271 | 301 | 10 | 5 |
| 3 | 2C, 4D-E, S7 | 167 | 15 | 7,271 | 301 | 10 | 5 |
| 4 | 2C, 4E, S7 | 168 | 15 | 7,271 | 301 | 10 | 5 |
| 5 | 2C, 4E, S7 | 169 | 15 | 7,271 | 301 | 10 | 5 |
| 6 | 2C, 4E, S7 | 170 | 15 | 7,271 | 301 | 10 | 5 |
| 7 | 2C, 4E, S7 | 171 | 15 | 8,703 | 301 | 10 | 6 |
| 8 | 2C, 4D-E, S7 | 172 | 15 | 8,703 | 301 | 10 | 6 |
| 9 | 2C, 4E, S7 | 173 | 15 | 8,703 | 301 | 10 | 6 |
| 10 | 2C, 4E, 6B, S7 | 174 | 15 | 8,703 | 301 | 10 | 6 |
| 11 | 2C, 4E, S7 | 175 | 15 | 8,703 | 301 | 10 | 6 |
| 12 | 2C, 4E, S7 | 176 | 15 | 8,703 | 301 | 10 | 6 |
| 13 | 2C, 4D-E, S7 | 177 | 15 | 8,703 | 301 | 10 | 6 |
| 14 | 2C, 4E, S7 | 178 | 15 | 8,703 | 301 | 10 | 6 |
| 15 | 2C, 4E, S7 | 179 | 15 | 10,135 | 301 | 10 | 7 |
| 16 | 2C, 4E, S7 | 180 | 15 | 10,135 | 301 | 10 | 7 |
| 17 | 2C, 4E, S7 | 181 | 15 | 10,135 | 301 | 10 | 7 |
| 18 | 2C, 4D-E, S7 | 182 | 15 | 10,135 | 301 | 10 | 7 |
| 19 | 2C, 4E, S7 | 183 | 15 | 10,135 | 301 | 10 | 7 |
| 20 | 2C, 4E, S7 | 184 | 15 | 10,135 | 301 | 10 | 7 |
| 21 | 2C, 4E, S7 | 185 | 15 | 10,135 | 301 | 10 | 7 |
| 22 | 2C, 4E, 7, S7 | 186 | 15 | 10,135 | 301 | 10 | 7 |
| 23 | 2C, 4D-E, S6, S7 | 187 | 15 | 11,567 | 301 | 10 | 8 |
| 24 | 2C, 4E, 6B, S7 | 188 | 15 | 11,567 | 301 | 10 | 8 |
| 25 | 2C, 4E, S7 | 189 | 15 | 11,567 | 301 | 10 | 8 |
| 26 | 2C, 4E, S7 | 190 | 15 | 11,567 | 301 | 10 | 8 |
| 27 | 2C, 4E, S7 | 191 | 15 | 11,567 | 301 | 10 | 8 |
| 28 | 2C, 4D-E, S7 | 192 | 15 | 11,567 | 301 | 10 | 8 |
| 29 | 2C, 4E, S7 | 193 | 15 | 11,567 | 301 | 10 | 8 |
| 30 | 2C, 4E, S7 | 194 | 15 | 11,567 | 301 | 10 | 8 |
| 31 | 2C, 4E, S7 | 195 | 15 | 13,000 | 301 | 10 | 9 |
| 32 | 2C, 4E, S7 | 196 | 15 | 13,000 | 301 | 10 | 9 |
| 33 | 2C, 4D-E, S7 | 197 | 15 | 13,000 | 301 | 10 | 9 |
| 34 | 2C, 4E, S7 | 198 | 15 | 13,000 | 301 | 10 | 9 |
| 35 | 2C, 4E, S7 | 199 | 15 | 13,000 | 301 | 10 | 9 |
| 36 | 2C, 4E, S7 | 200 | 15 | 13,000 | 301 | 10 | 9 |
| 37 | 2C, 4E, S7 | 201 | 15 | 13,000 | 301 | 10 | 9 |
| 38 | 2C, 4D-E, S7 | 202 | 15 | 13,000 | 301 | 10 | 9 |
| 39 | 2C, 4E, S7 | 203 | 15 | 14,432 | 301 | 10 | 10 |
| 40 | 2C, 4E, S7 | 204 | 15 | 14,432 | 301 | 10 | 10 |
| 41 | 2C, 4E, S7 | 205 | 15 | 14,432 | 301 | 10 | 10 |
| 42 | 2C, 4E, S7 | 206 | 15 | 14,432 | 301 | 10 | 10 |
| 43 | 2C, 4D-E, S7 | 207 | 15 | 14,432 | 301 | 10 | 10 |
| 44 | 2C, 4E, S7 | 208 | 15 | 14,432 | 301 | 10 | 10 |
| 45 | 2C, 4E, S7 | 209 | 15 | 14,432 | 301 | 10 | 10 |
| 46 | 2C, 4E, S7 | 210 | 15 | 14,432 | 301 | 10 | 10 |
| 47 | 2C, 4E, S7 | 211 | 15 | 15,864 | 301 | 10 | 11 |
| 48 | 2C, 4E, S7 | 212 | 15 | 15,864 | 301 | 10 | 11 |
| 49 | 2C, 4E, S7 | 213 | 15 | 15,864 | 301 | 10 | 11 |
| 50 | 2C, 4E, S7 | 214 | 15 | 15,864 | 301 | 10 | 11 |
| 51 | 2C, 4E, S7 | 215 | 15 | 15,864 | 301 | 10 | 11 |

* All included simulations are of dodecamers (12 nucleosomes) with 127 bp nucleosomal DNA wrapping.

**Supplementary Table S6:** Simulation parameters: dodecamer simulations with various nucleosomal DNA wrappings

| Index | Fig. Number | NRL (bp) | Inter-nucleosome Potential ($k_BT$) | Wrapping (bp) | Trials | Steps/ Snapshot | Total Snapshots | Burn-in | Beads/ Linker |
|---|---|---|---|---|---|---|---|---|---|
| 1 | 4C | 185 | 5.4 | 117 | 10 | 13,000 | 201 | 10 | 9 |
| 2 | 4B-C | 185 | 5.4 | 119 | 10 | 11,567 | 201 | 10 | 8 |
| 3 | 4C | 185 | 5.4 | 121 | 10 | 11,567 | 201 | 10 | 8 |
| 4 | 4B-C | 185 | 5.4 | 123 | 10 | 11,567 | 201 | 10 | 8 |
| 5 | 4C | 185 | 5.4 | 125 | 10 | 11,567 | 201 | 10 | 8 |
| 6 | 4B-C | 185 | 5.4 | 127 | 10 | 10,135 | 201 | 10 | 7 |
| 7 | 4C | 185 | 5.4 | 129 | 10 | 10,135 | 201 | 10 | 7 |
| 8 | 4B-C | 185 | 5.4 | 131 | 10 | 10,135 | 201 | 10 | 7 |
| 9 | 4C | 185 | 5.4 | 133 | 10 | 10,135 | 201 | 10 | 7 |
| 10 | 4B-C | 185 | 5.4 | 135 | 10 | 8,703 | 201 | 10 | 6 |
| 11 | 4C | 185 | 5.4 | 137 | 10 | 8,703 | 201 | 10 | 6 |
| 12 | 4B-C | 185 | 5.4 | 139 | 10 | 8,703 | 201 | 10 | 6 |
| 13 | 4C | 185 | 5.4 | 141 | 10 | 8,703 | 201 | 10 | 6 |
| 14 | 4B-C | 185 | 5.4 | 143 | 10 | 7,271 | 201 | 10 | 5 |
| 15 | 4C | 185 | 5.4 | 145 | 10 | 7,271 | 201 | 10 | 5 |
| 16 | 4B-C | 185 | 5.4 | 147 | 10 | 7,271 | 201 | 10 | 5 |
| 17 | S5 | 177 | 0 | 117 | 10 | 11,567 | 301 | 10 | 8 |
| 18 | S5 | 177 | 0 | 118 | 10 | 10,135 | 301 | 10 | 7 |
| 19 | S5 | 177 | 0 | 119 | 10 | 10,135 | 301 | 10 | 7 |
| 20 | S5 | 177 | 0 | 120 | 10 | 10,135 | 301 | 10 | 7 |
| 21 | S5 | 177 | 0 | 121 | 10 | 10,135 | 301 | 10 | 7 |
| 22 | S5 | 177 | 0 | 122 | 10 | 10,135 | 301 | 10 | 7 |
| 23 | S5 | 177 | 0 | 123 | 10 | 10,135 | 301 | 10 | 7 |
| 24 | S5 | 177 | 0 | 124 | 10 | 10,135 | 301 | 10 | 7 |
| 25 | S5 | 177 | 0 | 125 | 10 | 10,135 | 301 | 10 | 7 |
| 26 | S5 | 177 | 0 | 126 | 10 | 8,703 | 301 | 10 | 6 |
| 27 | S5 | 177 | 0 | 127 | 10 | 8,703 | 301 | 10 | 6 |
| 28 | S5 | 177 | 0 | 128 | 10 | 8,703 | 301 | 10 | 6 |
| 29 | S5 | 177 | 0 | 129 | 10 | 8,703 | 301 | 10 | 6 |
| 30 | S5 | 177 | 0 | 130 | 10 | 8,703 | 301 | 10 | 6 |
| 31 | S5 | 177 | 0 | 131 | 10 | 8,703 | 301 | 10 | 6 |
| 32 | S5 | 177 | 0 | 132 | 10 | 8,703 | 301 | 10 | 6 |
| 33 | S5 | 177 | 0 | 133 | 10 | 8,703 | 301 | 10 | 6 |
| 34 | S5 | 177 | 0 | 134 | 10 | 7,271 | 301 | 10 | 5 |
| 35 | S5 | 177 | 0 | 135 | 10 | 7,271 | 301 | 10 | 5 |
| 36 | S5 | 177 | 0 | 136 | 10 | 7,271 | 301 | 10 | 5 |
| 37 | S5 | 177 | 0 | 137 | 10 | 7,271 | 301 | 10 | 5 |
| 38 | S5 | 177 | 0 | 142 | 10 | 5,838 | 301 | 10 | 4 |
| 39 | S5 | 177 | 0 | 147 | 10 | 5,838 | 301 | 10 | 4 |

* All included simulations are of dodecamers (12 nucleosomes).

**Supplementary Table S7:** Simulation parameters: dodecamer simulations with various nucleosomal DNA wrappings and NRLs

| Index | Fig. Number | NRL (bp) | Wrapping (bp) | Trials | Steps/ Snapshot | Total Snapshots | Burn-in | Beads/Linker |
|---|---|---|---|---|---|---|---|---|
| 1 | 4F | 167 | 117 | 3 | 8,703 | 301 | 10 | 6 |
| 2 | 4F | 167 | 122 | 3 | 8,703 | 301 | 10 | 6 |
| 3 | 4F | 167 | 127 | 3 | 7,271 | 301 | 10 | 5 |
| 4 | 4F | 167 | 132 | 3 | 5,838 | 301 | 10 | 4 |
| 5 | 4F | 167 | 137 | 3 | 5,838 | 301 | 10 | 4 |
| 6 | 4F | 170 | 117 | 3 | 10,135 | 301 | 10 | 7 |
| 7 | 4F | 170 | 122 | 3 | 8,703 | 301 | 10 | 6 |
| 8 | 4F | 170 | 127 | 3 | 7,271 | 301 | 10 | 5 |
| 9 | 4F | 170 | 132 | 3 | 7,271 | 301 | 10 | 5 |
| 10 | 4F | 170 | 137 | 3 | 5,838 | 301 | 10 | 4 |
| 11 | 4F | 173 | 117 | 3 | 10,135 | 301 | 10 | 7 |
| 12 | 4F | 173 | 122 | 3 | 8,703 | 301 | 10 | 6 |
| 13 | 4F | 173 | 127 | 3 | 8,703 | 301 | 10 | 6 |
| 14 | 4F | 173 | 132 | 3 | 7,271 | 301 | 10 | 5 |
| 15 | 4F | 173 | 137 | 3 | 7,271 | 301 | 10 | 5 |
| 16 | 4F | 176 | 117 | 3 | 10,135 | 301 | 10 | 7 |
| 17 | 4F | 176 | 122 | 3 | 10,135 | 301 | 10 | 7 |
| 18 | 4F | 176 | 127 | 3 | 8,703 | 301 | 10 | 6 |
| 19 | 4F | 176 | 132 | 3 | 8,703 | 301 | 10 | 6 |
| 20 | 4F | 176 | 137 | 3 | 7,271 | 301 | 10 | 5 |
| 21 | 4F | 179 | 117 | 3 | 11,567 | 301 | 10 | 8 |
| 22 | 4F | 179 | 122 | 3 | 10,135 | 301 | 10 | 7 |
| 23 | 4F | 179 | 127 | 3 | 10,135 | 301 | 10 | 7 |
| 24 | 4F | 179 | 132 | 3 | 8,703 | 301 | 10 | 6 |
| 25 | 4F | 179 | 137 | 3 | 7,271 | 301 | 10 | 5 |
| 26 | 4F | 182 | 117 | 3 | 11,567 | 301 | 10 | 8 |
| 27 | 4F | 182 | 122 | 3 | 11,567 | 301 | 10 | 8 |
| 28 | 4F | 182 | 127 | 3 | 10,135 | 301 | 10 | 7 |
| 29 | 4F | 182 | 132 | 3 | 8,703 | 301 | 10 | 6 |
| 30 | 4F | 182 | 137 | 3 | 8,703 | 301 | 10 | 6 |
| 31 | 4F | 185 | 117 | 3 | 13,000 | 301 | 10 | 9 |
| 32 | 4F | 185 | 122 | 3 | 11,567 | 301 | 10 | 8 |
| 33 | 4F | 185 | 127 | 3 | 10,135 | 301 | 10 | 7 |
| 34 | 4F | 185 | 132 | 3 | 10,135 | 301 | 10 | 7 |
| 35 | 4F | 185 | 137 | 3 | 8,703 | 301 | 10 | 6 |
| 36 | 4F | 188 | 117 | 3 | 13,000 | 301 | 10 | 9 |
| 37 | 4F | 188 | 122 | 3 | 11,567 | 301 | 10 | 8 |
| 38 | 4F | 188 | 127 | 3 | 11,567 | 301 | 10 | 8 |
| 39 | 4F | 188 | 132 | 3 | 10,135 | 301 | 10 | 7 |
| 40 | 4F | 188 | 137 | 3 | 8,703 | 301 | 10 | 6 |
| 41 | 4F | 191 | 117 | 3 | 13,000 | 301 | 10 | 9 |
| 42 | 4F | 191 | 122 | 3 | 13,000 | 301 | 10 | 9 |
| 43 | 4F | 191 | 127 | 3 | 11,567 | 301 | 10 | 8 |
| 44 | 4F | 191 | 132 | 3 | 10,135 | 301 | 10 | 7 |
| 45 | 4F | 191 | 137 | 3 | 10,135 | 301 | 10 | 7 |
| 46 | 4F | 194 | 117 | 3 | 14,432 | 301 | 10 | 10 |
| 47 | 4F | 194 | 122 | 3 | 13,000 | 301 | 10 | 9 |
| 48 | 4F | 194 | 127 | 3 | 11,567 | 301 | 10 | 8 |
| 49 | 4F | 194 | 132 | 3 | 11,567 | 301 | 10 | 8 |
| 50 | 4F | 194 | 137 | 3 | 10,135 | 301 | 10 | 7 |

* All included simulations are of dodecamers (12 nucleosomes).

**Supplementary Table S7:** Simulation parameters: dodecamer simulations of various nucleosomal DNA wrappings and NRLs (continued)

| Index | Fig. Number | NRL (bp) | Wrapping (bp) | Trials | Steps/ Snapshot | Total Snapshots | Burn-in | Beads/Linker |
|---|---|---|---|---|---|---|---|---|
| 51 | 4F | 197 | 117 | 3 | 14,432 | 301 | 10 | 10 |
| 52 | 4F | 197 | 122 | 3 | 13,000 | 301 | 10 | 9 |
| 53 | 4F | 197 | 127 | 3 | 13,000 | 301 | 10 | 9 |
| 54 | 4F | 197 | 132 | 3 | 11,567 | 301 | 10 | 8 |
| 55 | 4F | 197 | 137 | 3 | 11,567 | 301 | 10 | 8 |
| 56 | 4F | 200 | 117 | 3 | 14,432 | 301 | 10 | 10 |
| 57 | 4F | 200 | 122 | 3 | 14,432 | 301 | 10 | 10 |
| 58 | 4F | 200 | 127 | 3 | 13,000 | 301 | 10 | 9 |
| 59 | 4F | 200 | 132 | 3 | 13,000 | 301 | 10 | 9 |
| 60 | 4F | 200 | 137 | 3 | 11,567 | 301 | 10 | 8 |
| 61 | 4F | 203 | 117 | 3 | 15,864 | 301 | 10 | 11 |
| 62 | 4F | 203 | 122 | 3 | 14,432 | 301 | 10 | 10 |
| 63 | 4F | 203 | 127 | 3 | 14,432 | 301 | 10 | 10 |
| 64 | 4F | 203 | 132 | 3 | 13,000 | 301 | 10 | 9 |
| 65 | 4F | 203 | 137 | 3 | 11,567 | 301 | 10 | 8 |
| 66 | 4F | 206 | 117 | 3 | 15,864 | 301 | 10 | 11 |
| 67 | 4F | 206 | 122 | 3 | 15,864 | 301 | 10 | 11 |
| 68 | 4F | 206 | 127 | 3 | 14,432 | 301 | 10 | 10 |
| 69 | 4F | 206 | 132 | 3 | 13,000 | 301 | 10 | 9 |
| 70 | 4F | 206 | 137 | 3 | 13,000 | 301 | 10 | 9 |
| 71 | 4F | 209 | 117 | 3 | 17,296 | 301 | 10 | 12 |
| 72 | 4F | 209 | 122 | 3 | 15,864 | 301 | 10 | 11 |
| 73 | 4F | 209 | 127 | 3 | 14,432 | 301 | 10 | 10 |
| 74 | 4F | 209 | 132 | 3 | 14,432 | 301 | 10 | 10 |
| 75 | 4F | 209 | 137 | 3 | 13,000 | 301 | 10 | 9 |
| 76 | 4F | 212 | 117 | 3 | 17,296 | 301 | 10 | 12 |
| 77 | 4F | 212 | 122 | 3 | 15,864 | 301 | 10 | 11 |
| 78 | 4F | 212 | 127 | 3 | 15,864 | 301 | 10 | 11 |
| 79 | 4F | 212 | 132 | 3 | 14,432 | 301 | 10 | 10 |
| 80 | 4F | 212 | 137 | 3 | 13,000 | 301 | 10 | 9 |

* All included simulations are of dodecamers (12 nucleosomes).

**Supplementary Table S8:** Simulation parameters: dodecamer simulations with heterogeneous linker lengths

| Index | Fig. Number | NRL (bp) | Trials | Steps/ Snapshot | Total Snapshots | Burn-in | Beads/Linker |
|---|---|---|---|---|---|---|---|
| 1 | 6A-B | 171$\pm$17 | 10 | 11,567 | 112 | 10 | 8 |
| 2 | 6A-B | 173$\pm$44 | 10 | 11,567 | 112 | 10 | 8 |
| 3 | 6A-B | 188$\pm$6 | 10 | 11,567 | 112 | 10 | 8 |
| 4 | 6A | 194$\pm$6 | 10 | 11,567 | 112 | 10 | 8 |
| 5 | 6A | 205$\pm$7 | 10 | 11,567 | 112 | 10 | 8 |
| 6 | 6A | 209$\pm$9 | 10 | 11,567 | 112 | 10 | 8 |

* All included simulations are of dodecamers (12 nucleosomes).

**Supplementary Table S9:** 1D-CNN model parameters and statistics

| Model Number | Description | Training Epochs | RMSE |
|---|---|---|---|
| 1 | Infer nucleosomal DNA wrapping | 10 | 1.01 bp |
| 2 | Infer nucleosomal DNA wrapping with ablated data | 10 | 1.23 bp |
| 3 | Infer NRL | 10 | 1.19 bp |
| 4 | Infer both NRL and nucleosomal DNA wrapping | 10 | 1.03 bp (wrap); 2.87 bp (NRL) |
| 5 | Infer inter-nucleosome strength | 100 | 0.563 $k_BT$ |
